# Functional-space alignment resolves the eco-evolutionary landscape of siderophore biosynthesis across bacteria

**DOI:** 10.1101/2025.03.09.642270

**Authors:** Jiqi Shao, Yanzhao Wu, Shizheng Tian, Ruichen Xu, Haohua Luo, Yuanzhe Shao, Linlong Yu, Ruolin He, Guanyue Xiong, Peng Guo, Nan Rong, Zhong Wei, Shaohua Gu, Zhiyuan Li

**Affiliations:** Quantitative Biology, Academy for Advanced Interdisciplinary Studies, Peking University, Beijing 100871, China; College of Resources and Environmental Sciences, State Key Laboratory of Nutrient Use and Management (SKL-NUM), National Academy of Agriculture Green Development, China Agricultural University, Beijing 100193, China; College of Computer Science and Technology (College of Data Science), Taiyuan University of Technology, Taiyuan, 030600, China; Peking-Tsinghua Center for Life Sciences, Academy for Advanced Interdisciplinary Studies, Peking University, Beijing 100871, China; Institute of ZooIogy, Chinese Academy of Sciences, Beijing 100101, China; Jiangsu Provincial Key Lab for Organic Solid Waste Utilization, Jiangsu Collaborative Innovation Center for Solid Organic Waste Resource Utilization, National Engineering Research Center for Organic-based Fertilizers, Nanjing Agricultural University, Nanjing 210095, China

## Abstract

Siderophores are central mediators of microbial iron acquisition, competition, and ecological adaptation, yet their biosynthetic diversity remains difficult to resolve across species because existing sequence-based BGC comparison is strongly constrained by phylogenetic background. Here we combine large-language-model-assisted literature mining, functional-space comparison, and genome-scale analysis to resolve the global organization of siderophore biosynthesis across bacteria. We first built SideroBank, a manually curated cross-species benchmark of siderophore biosynthetic gene clusters (BGCs), and used it to show that many identical products recur across distant taxa whereas the corresponding BGCs often fail to cluster in sequence space. We then developed BGC Block Aligner, which compares BGCs as ordered systems of functionally meaningful blocks and thereby converts comparison from sequence space to functional space. Applied to 97,432 bacterial genomes, this framework produced the Siderophore Atlas, revealing that siderophore synthesis is a remarkably pervasive trait encoded by over 60% of the analyzed genomes, with certain clusters being the most widely disseminated secondary metabolites across the bacterial domain. This global landscape suggests that the adoption of specific biosynthetic strategies is predominantly driven by ecological lifestyle rather than strict phylogenetic relatedness. Furthermore, a stark macro-evolutionary dichotomy **was** observed between the continuous structural diversification of NRPS pathways and the standardized, HGT-driven dissemination of NIS systems, linking functional-space genomics to the global ecology and evolution of siderophore biosynthesis..

## Introduction

Microbial natural products represent a major reservoir of chemical diversity and play central roles in survival, competition, and environmental adaptation. Advances in genome sequencing have transformed natural product discovery from a purely chemistry-driven enterprise into a comparative genomic problem, revealing immense and previously hidden biosynthetic potential across cultivated and uncultivated microorganisms^1^. Among natural products, siderophores are especially informative because they connect a chemically tractable biosynthetic system to a universal ecological problem: the acquisition of bioavailable iron. Beyond iron scavenging, siderophores also shape microbial competition, community interactions, host association, and pathogenicity, and have attracted growing attention in biotechnology and therapeutic development^2-4^. As such, resolving how siderophore biosynthesis is distributed across bacteria, and how similar products are encoded across divergent lineages, is central to understanding both microbial ecology and natural product evolution.

The rapid accumulation of genome and metagenome data has been accompanied by a broad suite of tools for BGC detection, annotation, and large-scale organization. antiSMASH has made genome-wide identification of biosynthetic pathways routine^5^, while DeepBGC and related machine-learning or deep-learning approaches have expanded BGC prediction and classification^6,7^. Curated resources such as MIBiG provide experimentally validated reference clusters for comparative analysis^8^. At the comparison stage, BiG-SCAPE and BiG-SLiCE have enabled large-scale clustering of BGCs by using domain content, domain order, and sequence similarity to define gene cluster families^9-11^, whereas pathway-specific tools such as RiPPMiner and TransATor demonstrate the value of incorporating biosynthetic logic for specialized metabolite classes^12,13^. Together, these frameworks have greatly accelerated the scalable exploration of biosynthetic diversity, but they mainly fundamentally rooted in sequence-based BGC comparison.

For many modular biosynthetic systems, however, sequence proximity is only an imperfect proxy for functional relatedness. In pathways such as NRPS and NRPS-independent(NIS), product identity is determined less by global full-length similarity than by the substrate specificity, functional identity, and ordered organization of key biosynthetic modules^14^. In NRPS systems, substrate choice is encoded in adenylation-domain recognition logic, classically captured by specificity-conferring residues and later extended by machine-learning-based predictors^15-18^. More broadly, NRPS diversification is shaped by recombination, exchange, and reorganization of modular units, emphasizing that local functional modules rather than full-length sequence similarity often determine biosynthetic output. In siderophore pathways, similar complications arise in NRPS-independent systems, where IucA/IucC-like synthetases and their active-site logic define core product assembly^3,19^. As a result, proteins from distant taxa may encode equivalent biosynthetic functions despite extensive sequence divergence, whereas globally similar proteins in closely related organisms may encode different substrate preferences. Functional differences and phylogenetic differences are therefore entangled in sequence space.

A second obstacle is that the reference knowledge needed to interpret siderophore diversity remains highly fragmented. Existing resources typically emphasize either chemical structures or a relatively small number of curated BGCs, whereas critical links among product identity, biosynthetic genes, and taxonomic distribution remain dispersed across the literature^8,20^. This fragmentation is particularly limiting for siderophores, whose biosynthesis has been investigated across many taxa, habitats, and research traditions but rarely organized into a unified cross-species reference system. Meanwhile, recent advances in protein structure prediction suggest that functionally constrained features may provide a more stable basis for cross-species comparison than full-length sequence similarity alone^21^. Together, these observations point to two unmet needs: a literature-derived cross-species benchmark that connects product identity to BGC architecture, and a comparison framework capable of recovering functional equivalence beyond the limits imposed by phylogenetic background.

Here, we address these limitations through an integrated framework that links literature-scale knowledge reconstruction to genome-scale functional comparison. We first developed Sidero-Mining, a large-language-model-assisted workflow for extracting siderophore biosynthetic knowledge from the literature, and used it to construct SideroBank, a manually curated cross-species benchmark of siderophore BGCs. Using this benchmark, we show that many siderophores recur across distant taxa whereas the corresponding BGCs often fail to cluster under existing sequence-based methods, revealing a major phylogenetic constraint in current comparison frameworks. We then developed BGC Block Aligner (BBA), which represents BGCs as ordered systems of functionally meaningful blocks and compares them through block-level alignment, thereby converting BGC comparison from sequence space to functional space. This framework enables functional typing of siderophore BGCs at a global scale, allowing us to build the Siderophore Atlas and to resolve the global organization, taxonomic spread, and contrasting macro-evolutionary regimes of NRPS and NIS siderophore pathways across bacteria.

## Result

### LLM-mined SideroBank exposes the phylogenetic constraint of sequence-based BGC comparison

High-quality reference datasets are essential for linking siderophore molecules to their biosynthetic gene clusters (BGCs), yet such resources remain incomplete. Existing databases either focus primarily on chemical structures or contain only limited numbers of annotated siderophore BGCs, whereas the key links among product identity, biosynthetic genes, and taxonomic distribution remain scattered across the literature. To address this gap, we developed Sidero-Mining, a large-language-model-assisted workflow for literature-scale extraction of siderophore biosynthetic information, and used it to construct SideroBank, a manually curated cross-species siderophore BGC resource (Figure 1A, Figure S1–S12). Through systematic screening of more than 10,000 siderophore-related publications, we retained 1,823 high-confidence articles for structured mining. After automated extraction, manual verification, genome retrieval, and BGC annotation, SideroBank compiled 738 nonredundant siderophore BGCs together with 325 NRPS adenylation-domain substrate annotations(Figure S13), substantially expanding the currently available siderophore reference space.

**Figure 1.**
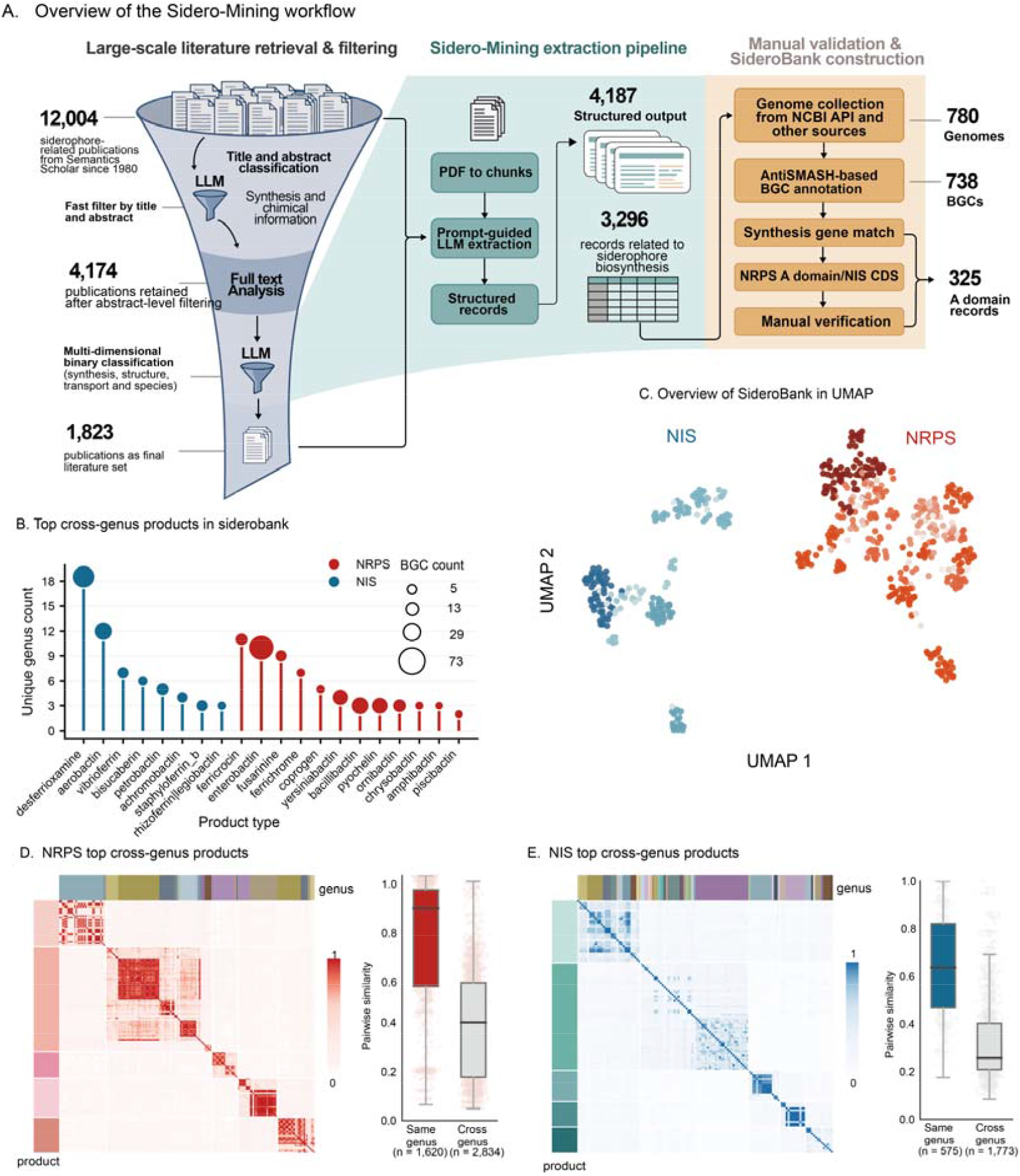
LLM-mined SideroBank reveals the phylogenetic constraint of sequence-based comparison for cross-species siderophore BGCs. (A) Schematic overview of the Sidero-Mining workflow. This pipeline uses a large language model for large-scale screening and structured extraction of siderophore-related literature, followed by manual curation, genome retrieval, and BGC annotation, ultimately generating SideroBank, a benchmark dataset of cross-species siderophore biosynthetic gene clusters. (B) Taxonomic distribution of the major cross-species siderophore products curated in SideroBank for NRPS and NIS pathways. (C) UMAP projection of pairwise sequence similarity among siderophore BGCs calculated by BiG-SCAPE. Red tones represent NRPS, and blue tones represent NIS. Color intensity indicates the abundance of BGCs within each product type. (D) Pairwise similarity heatmap and statistical analysis of cross-species same-product BGCs in the NRPS pathway. (E) Pairwise similarity heatmap and statistical analysis of cross-species same-product BGCs in the NIS pathway.

Systematic curation through SideroBank revealed that many siderophores recur across distantly related taxa, including across genera and higher taxonomic ranks, indicating that the taxonomic distribution of the same product is far broader than previously appreciated (Figure 1B). To further test whether existing comparison frameworks could reliably recognize these cross-species same-product BGCs, we used BiG-SCAPE to calculate pairwise similarity and visualize the resulting relationships in reduced dimensional space. For both NRPS and NIS pathways, BGCs derived from different species but encoding the same product often failed to cluster together in sequence similarity space (Figure 1C). Further analysis showed that similarity among BGCs encoding the same product was largely concentrated within genera, whereas pairwise similarity across genera was significantly lower. The mean within-genus versus between-genus similarity difference reached 0.46 for NRPS and 0.38 for NIS, indicating that substantial sequence divergence can arise even among cross-species same-product BGCs.

Because BiG-SCAPE measures similarity primarily from sequence similarity between matched domains, this sequence-space-based comparison is poorly suited for stably recognizing functionally equivalent BGCs across species and may systematically split them into separate groups. Overall, SideroBank not only revealed the widespread cross-species distribution of siderophore products, but also showed that existing sequence-based BGC comparison is insufficient to stably recover the correspondence among cross-species same-product BGCs. More importantly, by integrating a large number of manually curated cross-species same-product cases, SideroBank provides a critical benchmark dataset for evaluating the limitations of existing frameworks and for developing new methods capable of decoupling functional similarity from sequence divergence imposed by phylogenetic background.

### BGC Block Aligner converts BGC comparison from sequence space to functional space through block-level alignment

The inability of sequence-based BGC comparison to stably recover cross-species same-product BGCs indicates that global sequence similarity cannot serve as a reliable proxy for biosynthetic equivalence. Just as the biological meaning of nucleic acid and protein sequences is determined by the identity and order of their basic units, BGC function is likewise defined primarily by the functional identity, substrate specificity, and ordered organization of key biosynthetic modules. Based on this principle, we developed BGC Block Aligner (BBA). BBA first decomposes each BGC into functionally meaningful blocks, each representing a unit responsible for substrate recognition, scaffold construction, or key tailoring steps. These blocks are then organized into ordered block sequences according to their native biosynthetic organization and compared through block-level alignment, analogously to biological sequence alignment. In this way, BBA compares not raw sequences themselves, but function-defining units and their organizational relationships, thereby converting BGC comparison from sequence space to functional space(Figure 2A).

**Figure 2.**
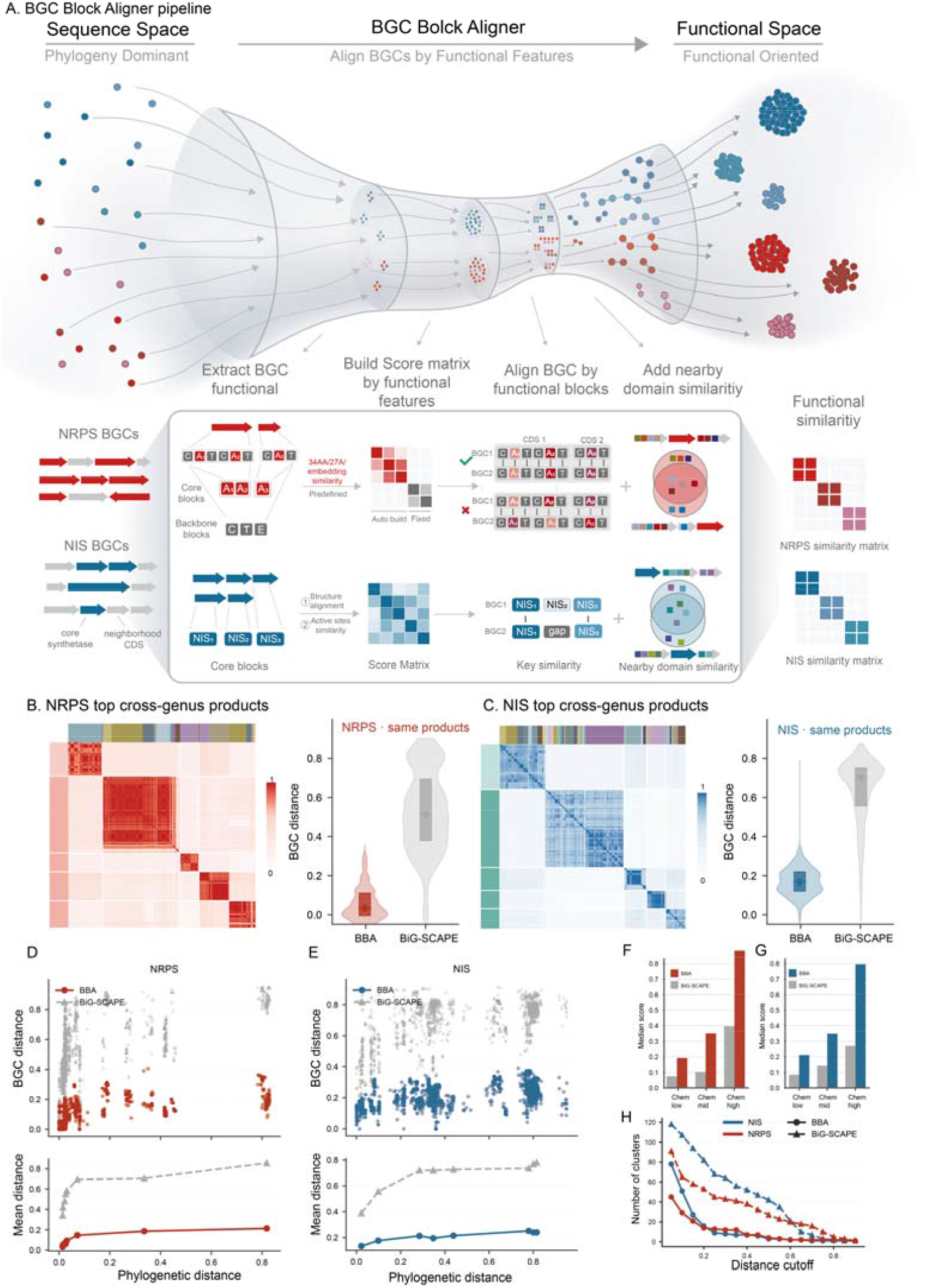
BGC Block Aligner converts siderophore BGC comparison from sequence space to functional space through block-level alignment. (A) Schematic overview of BGC Block Aligner (BBA). BBA first decomposes each BGC into functionally meaningful blocks, where each block corresponds to a functional unit involved in substrate recognition, scaffold construction, or key tailoring steps. For core blocks responsible for product scaffold formation, functional features are extracted to construct variable similarity matrices; for auxiliary backbone blocks, fixed scoring matrices are defined. These blocks are then organized into ordered block sequences according to their native biosynthetic arrangement and compared through block-level alignment. Similarity among nearby domains is further integrated to derive an overall functional similarity, thereby converting BGC comparison from sequence space to functional space. (B) Functional similarity matrix and clustering results for cross-species same-product BGCs in the NRPS pathway calculated by BBA. Compared with sequence-based methods, BBA more strongly organizes BGCs by product identity rather than by host taxonomy. (C) Functional similarity matrix and clustering results for cross-species same-product BGCs in the NIS pathway calculated by BBA. Similarly, BBA more stably recovers functional correspondence among cross-species same-product BGCs in the NIS pathway. (D) Relationship between pairwise BGC distance and host genome distance for NRPS same-product BGCs across species. The x-axis represents host genome distance and the y-axis represents pairwise BGC distance. Red circles indicate BBA, and grey triangles indicate BiG-SCAPE. (E) Relationship between pairwise BGC distance and host genome distance for NIS same-product BGCs across species. Blue circles indicate BBA, and grey triangles indicate BiG-SCAPE. (F) Relationship between BGC similarity and product chemical similarity in the NRPS dataset. Product pairs were grouped into three bins according to chemical similarity: Chem low (<0.3), Chem mid (0.3–0.7), and Chem high (>0.7). (G) Relationship between BGC similarity and product chemical similarity in the NIS dataset. As in NRPS, BBA-derived functional similarity shows stronger agreement with product chemical similarity in the NIS pathway. (H) Comparison of clustering results under different distance thresholds. Red and blue indicate NRPS and NIS siderophores, respectively; circles indicate BBA, and triangles indicate BiG-SCAPE.

**Figure 3.**
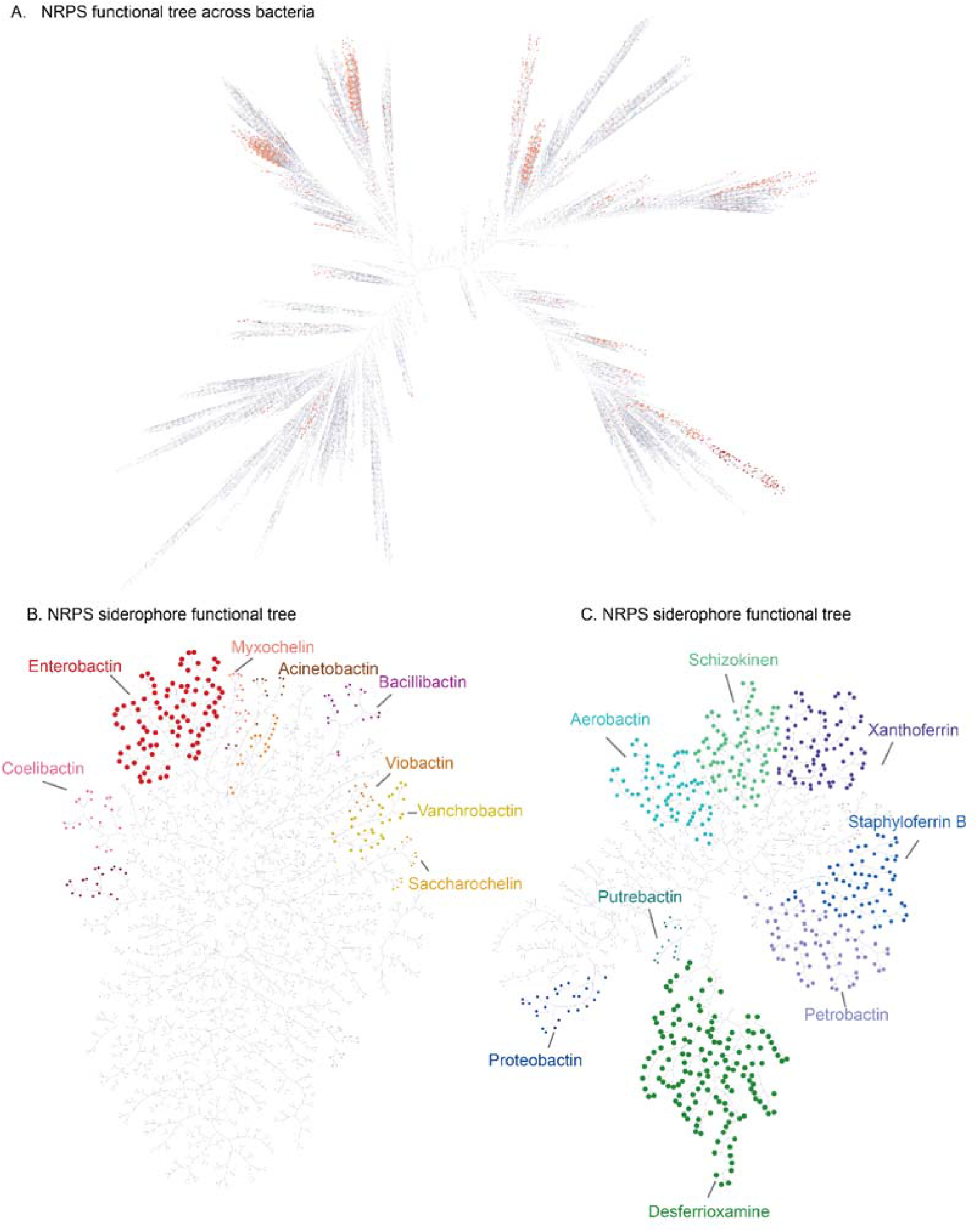
The Siderophore Atlas resolves the global functional organization of siderophore biosynthesis across bacterial genomes. (A) Global functional landscape of NRPS biosynthetic gene clusters across bacterial genomes. This map was generated by applying BGC Block Aligner to NRPS BGCs identified from 97,432 bacterial genomes. Each point represents a functionally typed NRPS BGC cluster observed in a given genus. Red tones indicate siderophore-associated NRPS clusters, and deeper color indicates broader cross-genus distribution. (B) Functional tree of more than 1,000 NRPS siderophore types. Each point represents a functionally typed NRPS BGC cluster observed in a given genus. Distinct colors highlight the top siderophore types showing the broadest cross-species distribution. (C) Functional tree of NIS siderophore types. Each point represents a functionally typed NIS BGC cluster observed in a given genus. Distinct colors highlight the top siderophore types showing the broadest cross-species distribution.

In BBA, block definition follows a function-first principle: core modules directly involved in product biosynthesis are extracted, whereas background genes unrelated to the target biosynthetic process are minimized; fragmented BGCs are merged using core domains as anchors when appropriate. For siderophore pathways, the most critical blocks are the biosynthetic modules that determine substrate recognition and core scaffold assembly, such as the A domain in NRPS pathways and IucA/IucC-like enzymes in NIS pathways. BBA further assigns block-level similarity matrices to different block types(Figure S14, S15). For NRPS, these matrices are based primarily on substrate-recognition features of A domains; for NIS, they incorporate structure-guided feature sequences derived from active-site-related positions. In both NRPS and NIS pathways, features extracted from substrate-recognition sites or structural constraints showed greater cross-species stability and stronger functional discrimination than full-length sequences, indicating that the signals most informative for cross-species comparison reside not in overall full-length similarity, but in function-defining sites directly related to biosynthetic activity.

We next evaluated BBA on the SideroBank benchmark dataset. Compared with BiG-SCAPE, the similarity matrices generated by BBA clustered BGCs more clearly by product identity rather than by host taxonomy, with the strong genus-level partitioning seen in sequence-based comparison markedly reduced (Figure 2B, C). In pairwise distance analyses, BBA produced substantially lower within-group distances for same-product BGCs than BiG-SCAPE: the mean distance was 0.09 for NRPS under BBA versus 0.50 under BiG-SCAPE, and 0.19 for NIS under BBA versus 0.80 under BiG-SCAPE, with markedly tighter distributions. When genome-level species distance was used as the x-axis to compare pairwise BGC distances across methods, BiG-SCAPE distances rose rapidly with increasing phylogenetic separation, whereas BBA functional distances increased only slightly and remained comparatively stable overall (Figure 2D, E), indicating much lower sensitivity to phylogenetic background. Moreover, when BGC similarity was stratified by product structural similarity, BBA showed substantially stronger agreement with chemical similarity than sequence-based comparison. In the subset with chemical similarity greater than 0.7, the mean functional similarity under BBA reached 0.78 for NRPS and 0.84 for NIS (Figure 2F, G). Under this metric, BBA also enabled more stable grouping at smaller thresholds and substantially reduced the inflation of cluster counts caused by phylogeny-driven sequence divergence (Figure 2H). Together, these results show that BBA provides a functional-space metric for cross-species BGC comparison that is more closely aligned with biosynthetic output than raw sequence similarity.

### Siderophore Atlas: high-resolution functional typing of siderophore BGCs in functional space

Having established the functional-space distance defined by BBA, we next applied it to large-scale functional typing of siderophore BGCs across bacterial genomes. Unlike conventional typing strategies that rely on sequence similarity, BBA provides a cross-species comparable measure of functional similarity, enabling a unified global view of siderophore biosynthetic potential. Guided by the distance distributions of manually curated same-product BGCs in SideroBank, we applied stringent criteria to perform fine-grained functional typing of NRPS and NIS BGCs and used the resulting classification to construct the Siderophore Atlas.

We applied BBA to 97,432 bacterial genomes and functionally typed 148,112 annotated NRPS BGCs, generating a global NRPS functional tree and identifying 7,479 distinct NRPS BGC functional types. Among these, 1,196 (15.1%) were assigned as dedicated NRPS-type siderophore biosynthetic types. Notably, although these siderophore-related functional types represent only a minority of all NRPS types, they were distributed across 48.3% of genomes, indicating striking ecological prevalence. These NRPS siderophore types were not confined to a small local region of the functional tree, but instead occupied multiple separated branches, and several types were broadly distributed across many genera. This pattern suggests that siderophore biosynthesis in NRPS functional space is not restricted to a single localized sector, but is supported by multiple differentiated combinations of function-defining modules.

Similarly, we functionally typed 39,246 NIS BGCs and identified 358 distinct NIS siderophore functional types. Together with the NRPS results, we ultimately defined 1,554 independent functionally defined BGC families for siderophore biosynthesis and named this global function-space-based classification framework the Siderophore Atlas. Overall, 64.8% of bacterial genomes encoded at least one siderophore biosynthetic system (Table S3), indicating that siderophore production is not restricted to a few specialized lineages but is instead a widespread adaptive capacity across bacterial genomes. Importantly, the Siderophore Atlas is not a catalog organized by conventional sequence clustering, but a high-resolution atlas organized by functional equivalence, enabling comparison of siderophore biosynthetic potential across phylogenetically diverse bacteria within a unified functional space.

### Macro-evolutionary landscape and taxonomic distribution of siderophore biosynthetic strategies

A global survey on siderophore prevalence across taxonomic levels reveals a paradoxical pattern. While siderophore biosynthesis is present in approximately two-thirds of genomes, its prevalence declines sharply at higher taxonomic ranks, dropping to ∼34% at the order level and ∼27% at the class level (Table S3). This steep reduction suggests that siderophore production is not universally conserved, but instead represents a highly selected trait shaped by specific ecological lifestyles.

To further resolve these lifestyle-associated pressures, we grouped bacterial classes based on their overall siderophore biosynthetic strategies (Figure 4A), revealing four distinct ecological archetypes that are also observed in other taxonomic levels (Figure S16):

1. The NRPS Generalists (archetype 1). Represented by Gammaproteobacteria and Actinomycetes, this archetype is characterized by a high abundance of NRPS siderophore BGCs and a frequent co-occurrence of NIS siderophores. These lineages invest heavily in complex modular biosynthetic pathways, consistent with adaptation to highly competitive and resource-limited microenvironments.
2. The NIS Specialists (archetype 2). Represented by classes such as Bacilli and Flavobacteriia, this group predominantly synthesize NIS siderophores while largely lacking canonical NRPS machinery. This streamlined strategy suggests specialization toward energetically efficient iron acquisition. Notably, although these two archetypes comprise less than half of all recognized bacterial classes, they encompass the majority of commonly sequenced taxa relevant to human health and agriculture, explaining their disproportionate representation at the genome level. For example, *Gammaproteobacteria* harbor notorious human pathogens and *Bacilli* have been widely utilized agricultural biocontrol agents, while *Alphaproteobacteria* and *Actinomycetes* are foundational to crop symbioses and soil biogeochemistry.
3. In contrast, the remaining classes reflect alternative survival strategies. The Non-siderophore NRPS Producers (archetype 3; spanning Desulfobacteria to Tissierellia) retain substantial NRPS capacity but appear to redirect these biosynthetic resources toward other classes of secondary metabolites (e.g., toxins, antimicrobials, or pigments) rather than siderophores. This reallocation is ecologically viable because these organisms secure iron through alternative avenues, such as anaerobic sulfate-reducers Desulfobacteria access soluble ferrous iron, and predatory lineages Myxococcia recycle iron from lysed prey, freeing their biosynthetic potential for chemical warfare and spatial competition.
4. The Minimalists (archetype 4; from Bacteroidia to Synergistia) lack both NRPS potential and siderophore biosynthetic pathways, consistent with a minimalists lifestyles such as obligate anaerobiosis or host dependence, where iron acquisition may rely on alternative mechanisms or iron-sparing strategies.

**Figure 4.**
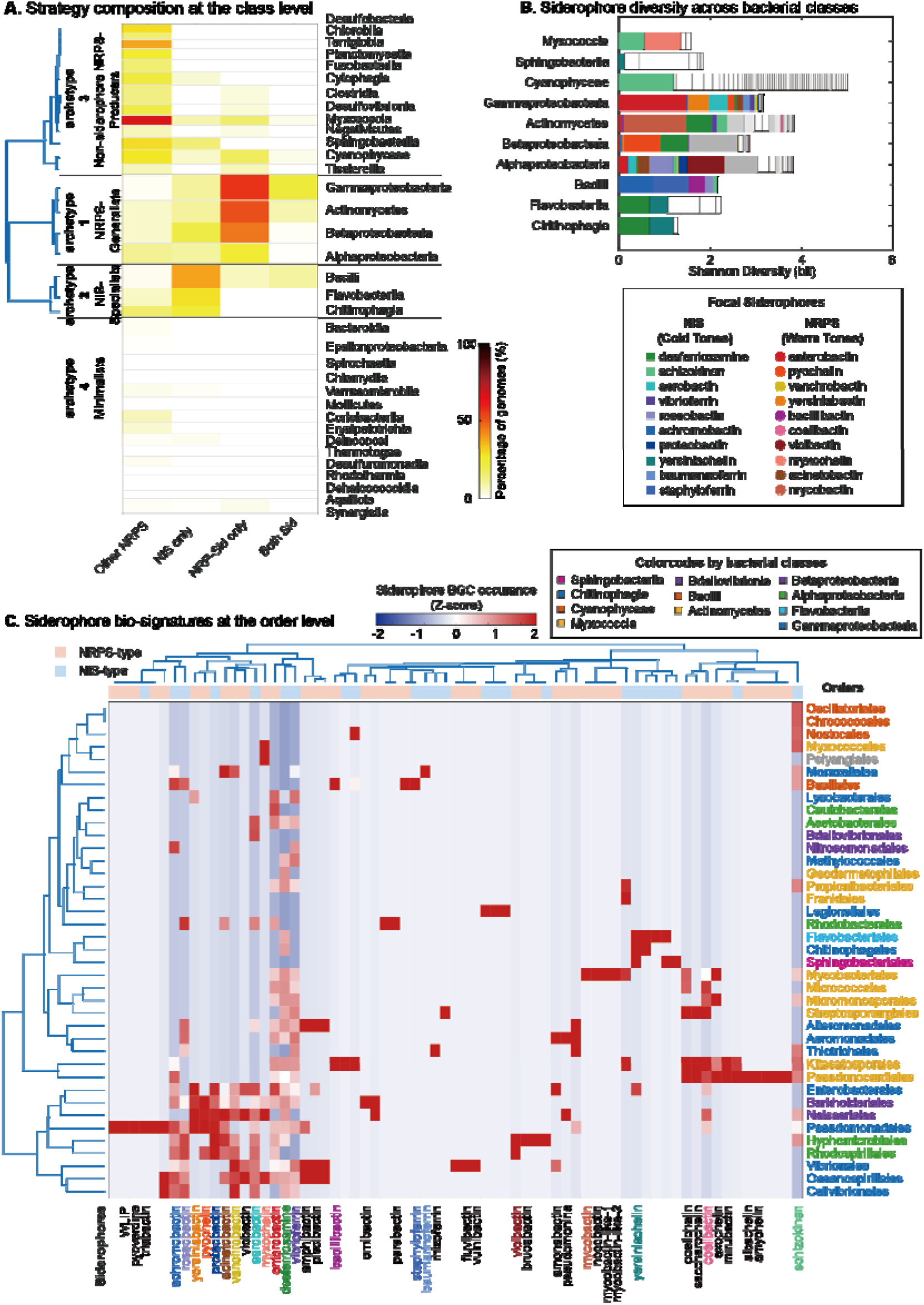
Macro-evolutionary landscape and taxonomic distribution of siderophore biosynthetic strategies. (A) Distribution of siderophore biosynthetic strategies across bacterial classes. The heatmap quantifies the percentage of genomes within each class adopting one of four mutually exclusive pathways: *Other NRPS* (possessing NRPS machinery but lacking dedicated siderophore BGCs); *NI-Sid only*; *NRP-Sid only*; and *Both Sid* (co-occurrence of both systems). Classes are vertically ordered by hierarchical clustering based on strategy profile similarity. (B) Compositional diversity of siderophore BGCs. The total length of each horizontal bar represents the Shannon Index (in bits), quantifying the cumulative siderophore diversity within each class. Bars are subdivided proportionally to reflect the relative abundance of specific BGCs. Focal siderophores are color-coded according to the legend (NIS in cold tones, NRPS in warm tones), while grey and white segments denote the minor contributions of other annotated and unannotated siderophore BGCs, respectively. (C) Biosignature landscape of siderophore potential across bacterial orders. A dual-hierarchical clustering heatmap illustrating the occurrence of distinct siderophore BGCs across varied orders. Color intensity denotes the Z-score normalized occurrence of each BGC (red: high; dark blue: absent/low). Rows represent bacterial orders, with labels color-coded by their parent class. Columns represent individual siderophore BGCs; labels for focal BGCs are colored consistently with plot B, non-focal BGCs with annotation are colored black, while unannotated BGCs are omitted for visual clarity. The top annotation band delineates the overarching biosynthetic logic: NRPS-type (peach) versus NIS-type (light blue). Clustering was performed using Ward’s linkage based on the Euclidean distance of prevalence profiles.

Focusing on bacterial classes with siderophore prevalence exceeding 10%, we observe substantial heterogeneity in their internal biosynthetic repertoires (Figure 4B). To characterize this variation, we tracked the proportional abundance of 20 focal siderophores representing major chemical classes of clinical and ecological significance. The resulting compositional profiles reveal a marked contrast in diversification strategies. Certain clades within the NIS Specialist archetype, such as Bacilli, exhibit relatively low internal diversity, dominated by a few highly successful focal siderophores. In contrast, lineages such as Actinomycetes display substantially broader biosynthetic diversity, driven largely by a long tail of non-focal and frequently unannotated BGCs. Accordingly, internal chemical diversity does not strictly correlate with overall siderophore prevalence, as evident in Cyanophyceae. These compositional differences warrant further investigation at the order level.

To further resolve these lifestyle-associated pressures at higher resolution, dual-hierarchical clustering of siderophore biosignatures (Figure 4C, Figure S17, Figure S 18) reveals that biosynthetic strategies across bacterial orders frequently group according to shared functional profiles, rather than phylogenetic relatedness. For example, certain orders within Gammaproteobacteria (e.g., Enterobacterales and Pseudomonadales) cluster in close proximity to Betaproteobacteria (e.g., Burkholderiales), suggesting convergence in siderophore repertoires. This pattern is consistent with adaptation to similarly competitive environments, where complex NRPS or hybrid siderophore systems may provide selective advantages. Conversely, marked divergence is observed even among closely related taxa. Within Actinomycetes, Mycobacteriales separates distinctly in the clustering space and is strongly dominated by mycobactin-like BGCs, whereas related orders such as Kitasatosporales and Pseudonocardiales display alternative repertoires, including NIS-type desferrioxamine and diverse NRPS pathways. This contrast highlights the functional plasticity of siderophore biosynthesis within phylogenetically related groups.

Shifting the focus from host taxonomy to the metabolites themselves, the heatmap reveals substantial variation in the distribution patterns of individual siderophores. Some BGCs, such as enterobactin and desferrioxamine, show relatively broad occurrence across multiple, phylogenetically distant orders, with recurring hotspots of high prevalence. This pattern is consistent with high evolutionary mobility and potential horizontal transfer. In contrast, other siderophores, including yersiniabactin and bacillibactin, exhibit more restricted distributions, largely confined to specific lineages. These differences point to distinct evolutionary trajectories, potentially reflecting variation in ecological specialization and dissemination capacity.

Together, these observations support a framework in which siderophore diversity is shaped not only by vertical inheritance but also by differential mobility and ecological selection, motivating a conceptual decoupling of siderophore repertoires from host taxonomy.

### Taxonomic spread reveals contrasting evolutionary patterns of NRPS and NIS pathways

The taxonomic distributions explored in the previous section expose a decoupling: a BGC’s prevalence at the genome level does not necessarily dictate its ability to permeate higher taxonomic ranks. To quantify this cross-lineage mobility, we introduce the metric of “taxonomic spread.” We define this as the absolute number of distinct host clades (e.g., classes) in which a specific BGC is detected, regardless of its genome frequency within those clades (Figure 5A). In essence, if a BGC appears in at least one genome of a given class, that class contributes to its total spread score.

**Figure 5.**
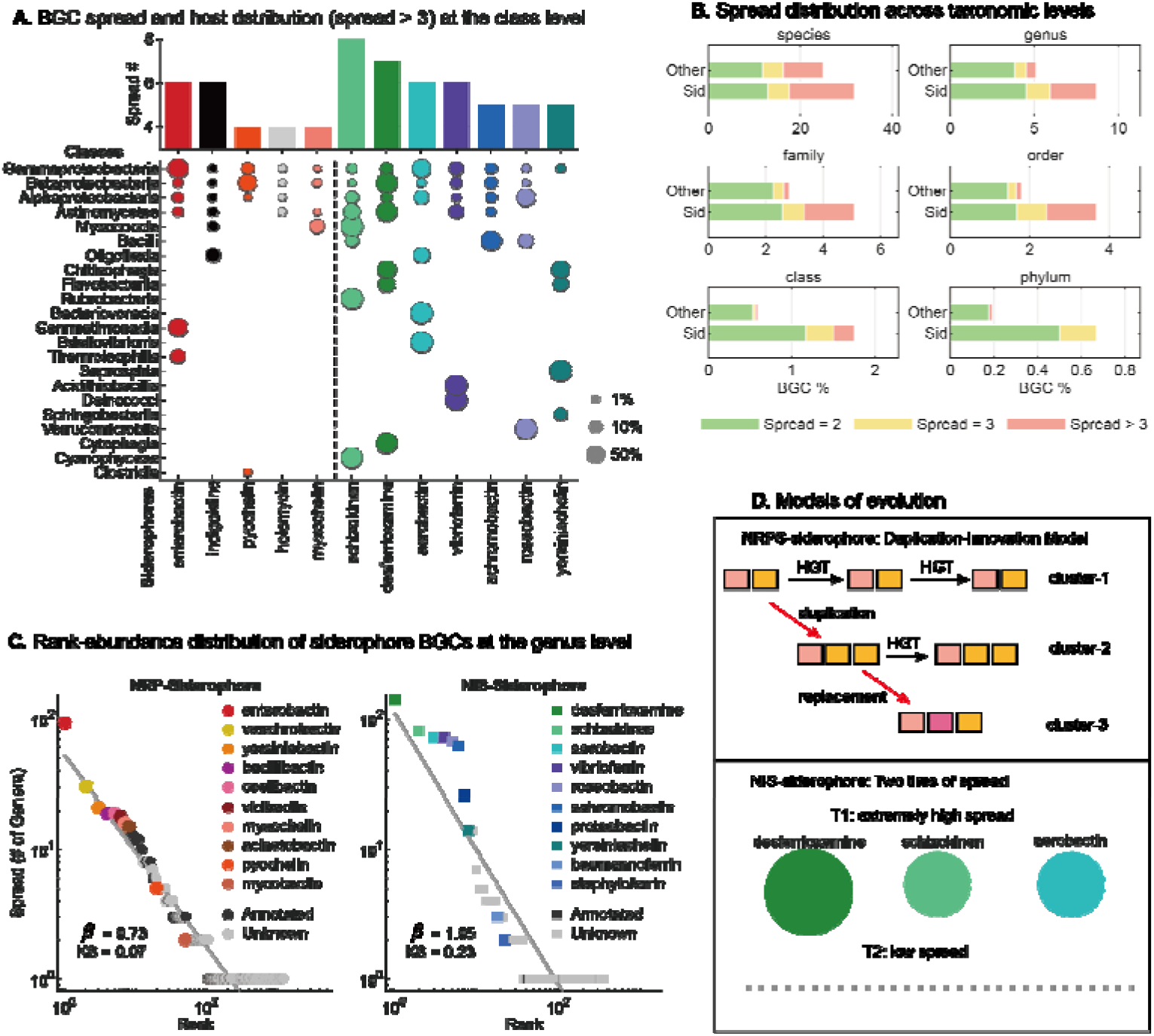
Taxonomic spread reveals contrasting evolutionary patterns of NRPS and NIS pathways. **(A) Taxonomic distribution of high-spread siderophore BGCs at the class level**. The top panel displays the specific spread value (number of distinct host classes) for each biosynthetic gene cluster (BGC), highlighting those with a spread greater than three. The lower bubble matrix illustrates the occurrence of these BGCs across host classes, where bubble size corresponds to the genome prevalence of a BGC within each taxon. “Star” or focal molecules are identified by their specific assigned colors (consistent with Figure 4), while other NRPS and NIS clusters are shown in black (e.g., indigoidine) and grey. Host taxa on the Y-axis are sorted by BGC richness, and BGCs on the X-axis are ordered by descending spread value. A vertical dashed line separates NRPS-type clusters (left) from NIS-type clusters (right). **(B) Comparative spread distribution of siderophore and non-siderophore NRPS BGCs across taxonomic levels**. The 3X2 panel illustrates the proportion of biosynthetic gene clusters (BGCs) with specific spread values (2, 3, and >3) relative to the total number of BGCs, Color coding distinguishes different spread values: light green (Spread=2), yellow (Spread = 3), and salmon (Spread > 3). “Sid” denotes siderophore-related NRPS clusters, and “Other” denotes non-siderophore NRPS clusters. Each subplot corresponds to a distinct taxonomic level, from species to phylum. **(C). Rank-abundance distribution of siderophore BGCs at the genus level**. The log-log plots illustrate the spread of NRP-siderophores (left panel) and NIS-siderophores (right panel) across host genera. Siderophores are ordered by their descending spread value on the X-axis (Rank), with the Y-axis representing the number of distinct host genera (Spread) for each BGC. The grey line indicates the Zipf (power-law) fit, with the resulting Zipf coefficient (beta) and Kolmogorov-Smirnov (KS) statistic provided in each panel. Focal molecules are highlighted with specific assigned colors, while other annotated and unknown clusters are shown in black and grey, respectively.

Visualizing the BGCs with the highest class-level spread (spread > 3; Figure 5A) highlights two preliminary trends. First, NRPS siderophores appear disproportionately represented among the most widely disseminated NRPS clusters. Despite comprising only 15% of the total NRPS repertoire, they account for three of the top five most broadly distributed NRPS BGCs, led by enterobactin (spread = 6). Second, the absolute highest spread values are dominated by a select group of NIS siderophores. Although NIS systems are present in a smaller fraction of global genomes (30%) compared to NRPS siderophores (∼48%), their top-ranking BGCs achieve exceptional cross-clade distribution. Seven distinct NIS pathways exceed a class-level spread of 3, spearheaded by schizokinen (spread = 8) and desferrioxamine (spread = 7).

Defined quantification confirmed that NRPS siderophores consistently exhibit a vastly superior expansion capacity compared to non-siderophore NRPSs. Specifically, the fraction of NRPS siderophore BGCs achieving broad cross-lineage distribution (i.e., successfully spanning >1, >2, or >3 distinct clades) substantially exceeds that of their non-siderophore counterparts, at taxonomic scales from genus to phylum. Complementary cumulative distribution function (CCDF) analysis confirms the statistical robustness of NPRS siderophores’s higher spread (Figure S19). Furthermore, from the species level up to the order level, the CCDF reveals that the spread frequencies for both NRPS categories follow a heavy-tailed distribution that tightly fits a power-law model, a scaling frequently carring intriguring eco-evolutionary implications.

To untangle the macro-ecological dynamics driving these pathways, we visualized the rank-abundance distributions of siderophore BGCs at the genus level (Figure 5C). Plotting taxonomic spread against rank on a log-log scale exposes fundamentally divergent evolutionary architectures between the two biosynthetic classes:

NRPS siderophores strictly adhere to a power-law distribution (KS = 0.07, *β* = 0.73). Scale-free is a typical character of such distribution: While canonical BGCs like enterobactin occupy the apex, the spread of subsequent clusters (such as vanchrobactin and yersiniabactin) decays log-linearly. This scale-free geometry precludes any defined threshold that scales the taxonomic spread, ultimately manifesting as a massive long tail of rare, frequently unannotated BGCs restricted to single genera. Ecologically, such power-law topologies can be classically explained by the Simon model, a generative “duplication-innovation” framework. Under this model, the empirical scaling exponent (*β* = 0.73) translates to an estimated innovation rate of approximately *u* =0.27. This remarkably high rate implies continuous, localized invention, perfectly mirroring the modular nature of NRPS assembly lines where domain swapping and recombination readily generate novel products (Figure 5D).

In contrast, the NIS siderophore distribution displays a cliff-like pattern. This deviation from power-law is confirmed statistically (*KS* = 0.23) and is visually apparent as a pronounced downward curvature on the log-log scale, culminating in a substantially steeper decay (*β* = 1.05) than that observed in NRPS. Unlike the continuous decay of rank-abundance in NRPS, spread of NIS clearly separates into two tiers: The top 6 or 8 highly mobilized focal pathways, including desferrioxamine, schizokinen, and aerobactin, attain massive taxonomic spread. This is then followed by a sharp decrease, where most pathways exhibit very low spread values less than 3(Figure 5C, gray dots). This mathematical profile, capped by a definitive diversity threshold, suggest an evolutionary reliance on horizontal gene transfer (HGT) of these “top-tier” solutions over *de novo* assembly-line invention.

Together, these contrasting mathematical profiles capture two fundamental evolutionary solutions to iron scarcity: the relentless structural diversification of NRPS systems versus the rapid, standardized dissemination of NIS pathways (Figure 5D).

## Discussion

Natural product genomics has transformed the discovery of microbial biosynthetic potential by shifting the field from culture-based compound isolation to large-scale genome- and metagenome-enabled analysis. Tools such as antiSMASH, BiG-SCAPE, and BiG-SLiCE have greatly improved BGC detection, annotation, and family-level organization, making comparative natural product genomics a central component of molecular discovery^5,9-11^. Yet our results indicate that, in the cross-species setting, sequence-based BGC comparison still faces a deeper bottleneck: global sequence similarity does not stably recover biosynthetic equivalence. This problem is particularly evident in siderophores, a class of natural products with well-defined molecular function, broad ecological importance, and strong clinical relevance. Siderophores are not only central to microbial iron acquisition but also participate in host-associated interactions, microbial competition, and therapeutic applications^2^. When cross-species same-product BGCs cannot be reliably recovered, the consequences extend beyond cluster classification to downstream functional annotation and ecological interpretation.

In this context, SideroBank is valuable not simply because it expands siderophore reference coverage, but because it establishes a missing cross-species benchmark dataset. Existing resources tend to emphasize either chemical structure, local cluster annotation, or a limited set of curated exemplars, but rarely connect product identity, BGC architecture, host distribution, and cross-species correspondence within one framework. By integrating a large number of manually curated cross-species same-product cases, SideroBank makes it possible to systematically ask whether BGCs associated with the same product should still be recognized as corresponding systems when they occur in distant lineages. Prior work has shown that siderophore pathways can be replaced, lost, or horizontally disseminated across bacterial groups, indicating that their distribution does not map trivially onto host phylogeny^22^. In this sense, SideroBank is not merely a resource but a reference coordinate system for evaluating the boundary conditions of cross-species BGC comparison.

Our study also highlights a deeper role for LLM-assisted knowledge extraction in natural product genomics. Here, the LLM does not merely accelerate literature mining; rather, it enables fragmented and heterogeneous biosynthetic knowledge to be formalized at scale. Sidero-Mining therefore acts as a knowledge reconstruction layer, converting literature-derived molecular knowledge into a structured reference space that can support benchmark construction, method development, and global functional typing. In other words, the LLM made knowledge reconstruction at scale possible, while manual curation made the resulting benchmark reliable. This suggests that the value of LLMs in natural product research may extend beyond annotation assistance to the construction of machine-readable biosynthetic reference systems.

Against this benchmark, BGC Block Aligner (BBA) addresses the central comparison problem at a more fundamental level by redefining what should be compared. Existing BGC comparison frameworks have greatly advanced natural product genomics, but they remain rooted largely in domain content, gene order, and full-length sequence similarity, making them better suited to recovering sequence proximity than stable functional equivalence. Prior work on NRPS and other modular assembly lines has shown that product diversification is closely linked to module exchange, local recombination, and the reorganization of substrate-selecting units^14^. Our results support the view that, for complex natural product BGCs, the key determinants of product scaffolds and functional divergence are not global full-length similarity, but the functional identity, substrate specificity, and ordered organization of core biosynthetic modules. Rather than treating a BGC as a contiguous stretch of raw sequence, BBA represents it as an ordered system of functionally meaningful blocks and compares these through block-level alignment, thereby moving BGC comparison from sequence space to functional space.

The significance of BBA extends beyond this conceptual shift. First, it provides a more biosynthetically grounded basis for defining functionally defined BGC families, reducing the tendency of phylogeny-driven sequence divergence to fragment closely related biosynthetic solutions into separate groups. Second, it improves the transfer of product-linked annotation into distant genomic backgrounds, making cross-species functional annotation more stable. Third, because BBA operates on functionally meaningful blocks, it provides a natural entry point for studying block-level replacement, recombination, insertion, deletion, and convergence in BGC evolution. At the same time, it helps distinguish true biosynthetic novelty from cases that appear novel only because sequence divergence obscures underlying functional similarity. In siderophores, this framework also provides an upstream foundation for linking biosynthesis to receptor specificity, resource sharing, and microbial interaction networks. With the emergence of structure prediction tools such as AlphaFold^21^, structural constraints and active-site features are increasingly becoming a more unified functional scale than full-length sequence similarity alone, consistent with our results for NIS pathways. The broader implication is that the signals most informative for cross-species comparison do not reside primarily in full-length similarity, but in substrate-recognition sites, active-site-associated features, and structurally constrained functional modules.

The resulting Siderophore Atlas extends this functional-space comparison to a global scale. Its value lies not in listing siderophore BGCs per se, but in organizing siderophore biosynthesis through functional typing rather than conventional sequence clustering. Our results show that, although siderophore biosynthesis is broadly prevalent across bacterial genomes, this prevalence does not imply simple conservation across higher taxonomic ranks. Instead, bacterial groups differ strikingly in their biosynthetic strategies, internal biosynthetic profiles, and taxonomic spread. This is consistent with broader views of siderophores as components of iron competition, host-associated interactions, and ecological networks, rather than merely as isolated iron chelators. Thus, the Siderophore Atlas is not a catalog organized by sequence clustering, but a global framework organized by functional equivalence, enabling siderophore biosynthetic potential to be compared across phylogenetically diverse bacteria on a unified scale.

At the macro-evolutionary level, our results further suggest that NRPS siderophores and NIS siderophores follow contrasting organizational regimes. NRPS siderophores exhibit a long-tailed, innovation-rich architecture, consistent with continuous diversification driven by modular recombination and local innovation. By contrast, NIS siderophores are dominated by a relatively small number of highly disseminated pathways, suggesting a dissemination-oriented architecture in which a few successful solutions account for much of the cross-lineage spread. This contrast is consistent with prior observations of recombination-driven NRPS diversification and siderophore pathway replacement or dissemination across bacterial lineages. In light of our taxonomic spread analysis, we interpret these patterns as evidence for at least two different biosynthetic macro-evolutionary strategies for coping with iron limitation: one based on continued expansion of chemical space through modular innovation, and another based on repeated dissemination of a limited set of highly transferable pathways. In this sense, siderophore diversity is not simply an accumulation of molecular variants, but a reflection of how different biosynthetic systems balance innovation against dissemination.

Several limitations should also be acknowledged. First, both SideroBank and the Siderophore Atlas remain constrained by currently sequenced genomes, available references, and annotation quality, and therefore cannot capture the full extent of siderophore diversity. Second, the functional equivalence recovered by BBA remains an inference based on biosynthetic features rather than direct chemical or genetic validation for every family. Third, BBA was developed and validated primarily in siderophore pathways; although its principles should be extensible to PKS and other modular natural product systems, the definition of blocks, extraction of functional features, and alignment rules will need to be adapted for each biosynthetic class. Even so, this study points to two broader methodological transitions that are now becoming feasible in natural product genomics: Sidero-Mining, aided by an LLM, converts fragmented biosynthetic knowledge in literature space into a computable reference space, whereas BBA further converts BGC comparison from a phylogeny-constrained sequence space into a more biosynthetically meaningful functional space. As structure prediction, substrate characterization, chemical annotation, and ecological datasets continue to improve, such a functional-space genomics framework may provide a more general route for linking gene clusters, molecular products, and community-level function.

## Methods

### Literature mining and construction of the SideroBank dataset

To construct a high-confidence cross-species siderophore knowledgebase, we first performed large-scale literature mining using the Semantic Scholar Dataset. By querying the Semantic Scholar API with the keyword *“siderophore”* and restricting publication years to post-1980 studies, we retrieved a total of 12,004 potentially relevant publications.

To evaluate the performance of different models in extracting siderophore-related information, we manually constructed a benchmark dataset consisting of 60 articles, including equal numbers of positive articles containing siderophore-related information and negative articles without such information. To reduce potential biases arising from publication period, specific siderophore names, or recurring cue words, the benchmark set was designed to include articles from different time periods and a diverse range of siderophore systems. Ground-truth labels were manually curated from these articles. Model outputs were then classified as true positives, false positives, true negatives, or false negatives, and overall precision, recall, and F1-score were calculated.

To improve the accuracy and consistency of literature extraction, we developed the Sidero-Mining pipeline with carefully designed task-specific prompts. Each prompt generally consisted of three components: (i) explicit specification of the extraction task, such as extracting siderophore names or producer species; (ii) a structured output template in Markdown format to facilitate downstream parsing, verification, and conversion into more visually organized formats such as XMind; and (iii) constraint instructions designed to suppress hallucinations, for example by requiring the model to return “null” for unavailable information and to avoid generating content outside the predefined template.

For siderophore biosynthesis extraction, we further implemented a multi-step reasoning workflow designed to mimic manual literature interpretation. In the first step, the model extracted all genes related to the target siderophore and summarized their putative biosynthetic functions. In the second step, these genes and their functional descriptions were used to infer the biosynthetic class of the siderophore. In the third step, the predicted biosynthetic type and the previously extracted gene set were jointly used to identify genes directly involved in siderophore synthesis. In the fourth step, the refined synthesis-gene information was combined with the original article content to extract substrate-level biosynthetic details. By decomposing a complex synthesis-mining task into a sequence of connected subtasks, this workflow enabled progressive reasoning and improved the accuracy of biosynthesis-related information extraction.

For large-scale screening, literature filtering was performed in two stages. In the first stage, titles and abstracts of the retrieved articles were input into gpt-4o-mini (gpt-4o-mini-2024-07-18), and six predefined prompts were used to classify articles into major topic categories, including siderophore chemical structure, biosynthesis, ecology, pathology, environmental and agricultural relevance, and metal-ion transport. For ease of downstream statistics, the model was instructed to return only binary “Y” or “N” outputs.

In the second stage, articles retained after title-and-abstract screening were further evaluated at the full-text level. Full texts were downloaded automatically using the links provided in the Semantic Scholar Dataset whenever possible, and the remaining articles were collected manually. Full texts were then processed using **gpt-4o-mini (gpt-4o-mini-2024-07-18)** with seven additional prompts to determine whether each article contained information related to single-gene studies, genomic studies, siderophore transport, iron-affinity measurements, siderophore receptors, chemical structure characterization, or microbial interactions. Articles primarily focused on siderophore synthesis and biosynthetic genes were retained as the final literature set for siderophore information mining.

### Genome retrieval and BGC annotation

After extracting siderophore-related information from the literature, we retrieved the corresponding genome data based on the strain information reported in the mined articles. To this end, we developed an automated genome retrieval workflow using the NCBI API. By querying strain names extracted from the literature, the program automatically downloaded the corresponding genome files from the NCBI Dataset. For strains that could not be directly matched due to renaming, taxonomic updates, or redundant strain-name annotations in the original articles, the relevant genome assemblies were collected manually from the NCBI website.

Once genome files were obtained, siderophore biosynthetic gene clusters were annotated using a custom annotation workflow based on antiSMASH 7.0 with default parameters. These annotations served as the primary basis for downstream BGC curation and comparison.

### Dataset collection and preprocessing

#### Manually curated cross-species BGC dataset

We first collected literature-reported records of siderophore distribution across species and assembled a cross-species reference dataset of siderophore biosynthetic gene clusters (BGCs). For each reported strain, the corresponding genome data were further examined and candidate BGCs were annotated. These annotations were then manually validated according to antiSMASH product-class assignments and the internal organizational structure of each BGC, to ensure consistency between the collected clusters and their reported siderophore products.

#### Predicted protein structure dataset

To establish a structural reference dataset for NIS biosynthetic enzymes, we queried InterPro using the Pfam entry corresponding to the IucA/IucC domain (IucA_IucC, PF04183) as the key domain and retrieved approximately 14,000 protein entries associated with AlphaFold2-predicted structures. Metadata for these entries, including the identifiers of their corresponding AlphaFold2 models, were obtained through the InterPro download API. The corresponding predicted protein structures were then downloaded through the AlphaFold database API.

Based on this dataset, amino acid sequences were extracted and compiled into a searchable DIAMOND sequence database, which was used in subsequent analyses to match query NIS biosynthetic enzymes to reference sequences and structures.

### BGC Block Aligner algorithm and parameter settings

#### Definition of core biosynthetic domains

For NRPS pathways, the set of domains used for BGC comparison was defined based on the antiSMASH annotation framework for NRPS core components. This set included both the major catalytic domain subtypes that constitute canonical NRPS modules and additional domains involved in modification or accessory biosynthetic functions. These domain definitions were used for downstream module decomposition, feature extraction, and scoring-matrix construction.

For NIS pathways, the analysis focused on the key NIS biosynthetic enzymes responsible for siderophore assembly, which served as the basic units for feature extraction and similarity comparison.

#### Extraction features from NRPS A domains

Because adenylation (A) domains are the primary determinants of substrate specificity in NRPS pathways, we extracted multiple types of sequence-derived features from A domains for downstream similarity calculation. First, the 34-amino-acid (34AA) active-site feature sequences were extracted using a modified implementation based on the source code of AmenPredictor. In this procedure, protein sequences were aligned against a given hidden Markov model using HMMER2, and residues corresponding to predefined active-site positions in the template were extracted as feature residues.

In addition, we extracted the 27-amino-acid (27AA) specificity-conferring features defined in DeepAden, enabling an alternative representation of the substrate-determining region of each A domain. To further capture broader contextual information beyond manually defined active-site residues, we also obtained embedding features for each A domain using the DeepAden framework. These embeddings provide a continuous representation of sequence-level functional properties and were used as an additional feature space for pairwise comparison.

After feature extraction from all A domains in the BGCs to be compared, redundant feature sequences were removed using DIAMOND or CD-HIT where applicable. Pairwise similarity between A domains was then quantified on the basis of sequence identity between the extracted 34AA or 27AA feature sequences, or by similarity measures computed from the corresponding embedding representations. These similarity values were subsequently used for scoring-matrix construction.

#### Feature extraction for NIS biosynthetic enzymes

For NIS pathways, we first constructed a DIAMOND sequence database from the predicted protein structure dataset described above. Query NIS biosynthetic enzyme sequences were searched against this database, and the highest-scoring hit passing the predefined threshold was selected and mapped to its corresponding AlphaFold2-predicted structure. When no related sequence could be identified under the preset threshold of 0.9, the query structure was predicted using ESMFold.

After obtaining the predicted structures of the query NIS enzymes, pairwise structural alignments were performed between query and reference proteins using gtalign. Based on the active-site positions defined in the reference templates, the corresponding residues in the aligned query sequences were extracted as active-site feature sequences for downstream similarity calculation.

Different reference proteins were used for different classes of NIS biosynthetic enzymes, including DesD (PDB: 7tgm), IucA (PDB: 5jm8), SbnC (PDB: 7cbb), and AsbB (PDB: 3to3). For each reference, the active-site residue positions were defined according to the corresponding structural template and used to extract the aligned residues from the query proteins.

#### Construction of the scoring matrices

For NRPS pathways, a unified scoring matrix was constructed by combining A domains with other biosynthetically relevant domains. For non-A domains, a match between domains of the same family and the same subtype was assigned a score of 0.2, whereas a match between domains of the same family but different subtypes was assigned a score of 0.12. A mismatch between domains from different families was penalized by -0.1. Because A domains carry the most important functional information, their match scores were not assigned as fixed constants but were instead derived directly from the pairwise similarity matrix of A-domain feature sequences, or from transformed values based on that matrix, with scores constrained to the range of 0 to 1. To prevent biologically implausible matches between A domains and non-A domains, such matches were penalized by -0.5. The final NRPS scoring matrix therefore integrated both A-domain similarities and scores for other biosynthetic domains.

For NIS pathways, the relative order of biosynthetic enzymes was not explicitly considered. Instead, the analysis focused on similarity relationships among NIS enzymes within each pair of BGCs. Active-site feature sequences extracted from each structural template were used to generate separate similarity matrices, and for each enzyme pair, the maximum similarity value across these matrices was retained as the final similarity score. This final similarity matrix, with values ranging from 0 to 1, was then used directly, or after optional transformation, as the scoring matrix for downstream optimization.

#### Generation of pathway arrangements and BGC comparison

For NRPS pathways, all NRPS modules were treated as the basic units for arrangement and comparison. Under the exhaustive arrangement mode, when the number of module units was below a predefined threshold, such as five, all possible permutations of the modules were generated and evaluated. Considering the linear biosynthetic logic of NRPS systems, adjacent modules located within the same operon and oriented in the same transcriptional direction could be merged into a single unit, thereby reducing the size of the permutation space and minimizing fragmentation of contiguous module blocks. When the number of module units exceeded the threshold, a greedy strategy was used instead. Specifically, the BGC containing the largest number of modules was selected as the reference BGC, and its module units were fixed according to their genomic order. As in the exhaustive mode, adjacent and co-directional modules within the same operon could be merged into a single unit. For each query BGC, all module units were extracted and locally scored against different regions of the reference arrangement using the predefined scoring matrix. Candidate segments were then iteratively placed according to their local scores, and penalties were applied when newly placed segments overlapped with regions that had already been assigned in previous steps.

After the arrangement step, overall NRPS BGC comparison was performed on the concatenated domain sequences using the Biopython pairwise aligner under the specified scoring matrix. In the exhaustive arrangement mode, all possible arrangements were evaluated and the highest-scoring alignment was retained as the final result. In the greedy mode, the alignment score of the assembled arrangement was used as the final similarity score for the BGC pair.

For NIS pathways, no direct global ordering of biosynthetic enzymes was imposed. Instead, the method evaluated the quality of correspondence between the sets of NIS enzymes in the two BGCs. Based on the pairwise similarity scoring matrix described above, the Hungarian algorithm was used to identify the optimal enzyme matching between the two BGCs. The resulting total matching score was then normalized by the larger number of NIS enzymes present in either BGC, thereby reducing the effect of copy-number differences on the final similarity estimate.

#### BiG-SCAPE parameter settings

To benchmark the results of BGC Aligner against an established BGC comparison framework, the same set of siderophore BGCs was also analyzed using BiG-SCAPE. BiG-SCAPE was run with Pfam version 34, using the --no_classify option and --mix mode. Computations were performed with 32 CPU cores, and the cutoff was set to 1 to retain all pairwise comparison results. All other parameters were kept at their default settings. The raw similarity values output by BiG-SCAPE were extracted and further assembled into complete pairwise similarity matrices for downstream analyses.

## Author contributions

Jiqi Shao: Conceptualization; data curation; algorithm development; analysis; visualization; writing - original draft. Yanzhao Wu: Data curation; visualization; coding and analysis; writing - original draft. Shizhen Tian: Data curation. Haohua Luo: Data curation. Yuanzhe Shao: Data curation. Linlong Yu: Data curation. Guanyue Xiong:Data curation. Ruolin He: Data curation. Ruichen Xu: Data curation. Peng Guo: Data curation. Nan Rong:Data curation. Shaohua Gu: Conceptualization; funding acquisition. Zhiyuan Li: Conceptualization; funding acquisition; analysis; visualization; writing - original draft.

## Reference

1. Medema, M.H., and Fischbach, M.A. (2015). Computational approaches to natural product discovery. Nat Chem Biol 11, 639–648. 10.1038/nchembio.1884.

2. Kramer, J., Özkaya, Ö., and Kümmerli, R. (2020). Bacterial siderophores in community and host interactions. Nature Reviews Microbiology 18, 152–163. 10.1038/s41579-019-0284-4.

3. Carroll, C.S., and Moore, M.M. (2018). Ironing out siderophore biosynthesis: a review of non-ribosomal peptide synthetase (NRPS)-independent siderophore synthetases. Crit Rev Biochem Mol Biol 53, 356–381. 10.1080/10409238.2018.1476449.

4. Wei, Z., Gu, S., Vollenweider, V., Zuo, Y., Li, Z., and Kümmerli, R. (2025). Microbial siderophores for One Health. Trends in Microbiology 33, 1277–1285. 10.1016/j.tim.2025.05.002.

5. Blin, K., Shaw, S., Augustijn, H.E., Reitz, Z.L., Biermann, F., Alanjary, M., Fetter, A., Terlouw, B.R., Metcalf, W.W., Helfrich, E.J.N., et al. (2023). antiSMASH 7.0: new and improved predictions for detection, regulation, chemical structures and visualisation. Nucleic Acids Research 51, W46–W50. 10.1093/nar/gkad344.

6. Hannigan, G.D., Prihoda, D., Palicka, A., Soukup, J., Klempir, O., Rampula, L., Durcak, J., Wurst, M., Kotowski, J., Chang, D., et al. (2019). A deep learning genome-mining strategy for biosynthetic gene cluster prediction. Nucleic Acids Research 47, e110–e110. 10.1093/nar/gkz654.

7. Lai, Q., Yao, S., Zha, Y., Zhang, H., Zhang, H., Ye, Y., Zhang, Y., Bai, H., and Ning, K. (2025). Deciphering the biosynthetic potential of microbial genomes using a BGC language processing neural network model. Nucleic Acids Research 53, gkaf305. 10.1093/nar/gkaf305.

8. Zdouc, Mitja M., Blin, K., Louwen, Nico L.L., Navarro, J., Loureiro, C., Bader Chantal D., Bailey, Constance B., Barra, L., Booth, Thomas J., Bozhüyük, Kenan A.J., et al. (2024). MIBiG 4.0: advancing biosynthetic gene cluster curation through global collaboration. Nucleic Acids Research 53, D678–D690. 10.1093/nar/gkae1115.

9. Kautsar, S.A., van der Hooft, J.J.J., de Ridder, D., and Medema, M.H. (2021). BiG-SLiCE: A highly scalable tool maps the diversity of 1.2 million biosynthetic gene clusters. GigaScience 10, giaa154. 10.1093/gigascience/giaa154.

10. Navarro-Muñoz, J.C., Selem-Mojica, N., Mullowney, M.W., Kautsar, S.A., Tryon, J.H., Parkinson, E.I., De Los Santos, E.L.C., Yeong, M., Cruz-Morales, P., Abubucker, S., et al. (2020). A computational framework to explore large-scale biosynthetic diversity. Nature Chemical Biology 16, 60–68. 10.1038/s41589-019-0400-9.

11. Draisma, A., Loureiro, C., Louwen, N.L.L., Kautsar, S.A., Navarro-Muñoz, J.C., Doering, D.T., Mouncey, N.J., and Medema, M.H. (2026). BiG-SCAPE 2.0 and BiG-SLiCE 2.0: scalable, accurate and interactive sequence clustering of metabolic gene clusters. Nature Communications 17, 2000. 10.1038/s41467-026-68733-5.

12. Agrawal, P., Khater, S., Gupta, M., Sain, N., and Mohanty, D. (2017). RiPPMiner: a bioinformatics resource for deciphering chemical structures of RiPPs based on prediction of cleavage and cross-links. Nucleic Acids Research 45, W80–W88. 10.1093/nar/gkx408.

13. Helfrich, E.J.N., Ueoka, R., Dolev, A., Rust, M., Meoded, R.A., Bhushan, A., Califano, G., Costa, R., Gugger, M., Steinbeck, C., et al. (2019). Automated structure prediction of trans-acyltransferase polyketide synthase products. Nat Chem Biol 15, 813–821. 10.1038/s41589-019-0313-7.

14. Calcott, M.J., Owen, J.G., and Ackerley, D.F. (2020). Efficient rational modification of non-ribosomal peptides by adenylation domain substitution. Nat Commun 11, 4554. 10.1038/s41467-020-18365-0.

15. Stachelhaus, T., Mootz, H.D., and Marahiel, M.A. (1999). The specificity-conferring code of adenylation domains in nonribosomal peptide synthetases. Chem Biol 6, 493–505. 10.1016/s1074-5521(99)80082-9.

16. Röttig, M., Medema, M.H., Blin, K., Weber, T., Rausch, C., and Kohlbacher, O. (2011). NRPSpredictor2—a web server for predicting NRPS adenylation domain specificity. Nucleic Acids Research 39, W362–W367. 10.1093/nar/gkr323.

17. Rausch, C., Weber, T., Kohlbacher, O., Wohlleben, W., and Huson, D.H. (2005). Specificity prediction of adenylation domains in nonribosomal peptide synthetases (NRPS) using transductive support vector machines (TSVMs). Nucleic Acids Res 33, 5799–5808. 10.1093/nar/gki885.

18. Mongia, M., Baral, R., Adduri, A., Yan, D., Liu, Y., Bian, Y., Kim, P., Behsaz, B., and Mohimani, H. (2023). AdenPredictor: accurate prediction of the adenylation domain specificity of nonribosomal peptide biosynthetic gene clusters in microbial genomes. Bioinformatics 39, i40–i46. 10.1093/bioinformatics/btad235.

19. Yang, J., Banas, V.S., Patel, K.D., Rivera, G.S.M., Mydy, L.S., Gulick, A.M., and Wencewicz, T.A. (2022). An acyl-adenylate mimic reveals the structural basis for substrate recognition by the iterative siderophore synthetase DesD. Journal of Biological Chemistry 298. 10.1016/j.jbc.2022.102166.

20. He, R., Gu, S., Xu, J., Li, X., Chen, H., Shao, Z., Wang, F., Shao, J., Yin, W.-B., Qian, L., et al. (2024). SIDERITE: Unveiling hidden siderophore diversity in the chemical space through digital exploration. iMeta 3, e192. 10.1002/imt2.192.

21. Jumper, J., Evans, R., Pritzel, A., Green, T., Figurnov, M., Ronneberger, O., Tunyasuvunakool, K., Bates, R., Žídek, A., Potapenko, A., et al. (2021). Highly accurate protein structure prediction with AlphaFold. Nature 596, 583–589. 10.1038/s41586-021-03819-2.

22. Bruns, H., Crüsemann, M., Letzel, A.C., Alanjary, M., McInerney, J.O., Jensen, P.R., Schulz, S., Moore, B.S., and Ziemert, N. (2018). Function-related replacement of bacterial siderophore pathways. Isme j 12, 320–329. 10.1038/ismej.2017.137.

