## Supplementary Information for "Functional-space alignment resolves the eco-evolutionary landscape of siderophore biosynthesis across bacteria"

### Macro-evolutionary Dynamics and the Simon Model

1. **The Simon Model: Core Process and Evolutionary Analogy**

The Simon model is a stochastic generative process that yields the Yule–Simon distribution in growing systems. Models of this class have been widely used in evolutionary and ecological contexts to explain heavy-tailed distributions, including species abundance and gene family size distributions. These applications rely on the same underlying principle of innovation coupled with preferential expansion, making the framework directly analogous to the diversification and spread of BGC clusters considered in this work.

**1.1 Defining the Evolutionary "Event"**

The Simon process is discrete. We define a single “event” ($t \to t+1$) as the addition of one instance to the total population—specifically, one BGC cluster appearing in a new taxonomic clade (e.g., a genus, order, or phylum). At each step, there are two mutually exclusive possibilities for this new occurrence:

1. **Innovation (Probability** $\mu$**):** With a constant probability $\mu$, the new instance represents a **novel BGC cluster**. In NRPS, this is formed by radical module recombination, domain swapping, or de novo assembly.
2. **Duplication/Preferential Attachment (Probability** $1-\mu$**):** With probability $1-\mu$, the new instance is an **expansion of an existing cluster**. Crucially, the probability that a specific cluster $i$ is chosen is proportional to its current “success”, defined by its taxonomic spread $k_{i}$ (the number of taxa clades it already occupies).
   1.2. **Analogy to NRPS Evolution**

In the context of NRPS macro-ecology, we define:

- **Categories:** Distinct NRPS clusters (e.g., the enterobactin cluster, the yersiniabactin cluster, etc).
- **Category Size (**$k$**):** The **Taxonomic Spread**, representing the number of distinct taxonomic clades (at a given level) a NRPS cluster has colonized.
- **Innovation Rate (**$\mu$**):** The frequency at which domain swapping or module recombination in NRPS creating a “novel” cluster that begins its own expansion history.
- **Duplication/Preferential Attachment:** A BGC cluster that is already widely distributed has more templates available for further dissemination, resembling the “rich-get-richer” mechanism.
  - **Mechanisms of Expansion:** Expansion occurs via **Vertical Transmission (VT)** or **Horizontal Gene Transfer (HGT)**.
  - **Scale-Dependency:** At lower taxonomic levels (e.g., within the same genus), VT (speciation) and HGT are highly frequent, leading to high duplication rates ($1-\mu$) and lower $\mu$. However, at higher levels (e.g., order level), VT is ineffective for crossing clade boundaries, and inter-order HGT faces immense biological and ecological barriers. Consequently, as the taxonomic scale increases, the relative probability of duplication decreases, leading to an effectively higher **innovation rate (**$\mu$**)**.

**2. Master Equation Derivation**

Let $N\left( t \right)$ be the total number of BGC occurrences across all clades at step $t$. Since one occurrence is added per step, $N\left( t \right)=t$ (ignoring initial conditions in the large-$t$ limit). Let $m_{k}\left( t \right)$ be the expected number of BGC clusters having a taxonomic spread of exactly $k$ at time $t$.

**2.1 The Master Equations**

The change in the number of BGC clusters with spread $k$ between step $t$ and $t+1$ is governed by the following transitions:

**For** $k > 1$**:** A cluster moves into the “spread $k$” bin if a cluster of spread $k-1$ expands. A family leaves the “spread $k$” bin if it expands to $k+1$.

$$\Delta m_{k}=m_{k}\left( t+1 \right)-m_{k}\left( t \right)=\left( 1-\mu\right)\frac{\left( k-1 \right)m_{k-1}\left( t \right)}{t}-\left( 1-\mu\right)\frac{km_{k}\left( t \right)}{t}.\quad$$

Eq S1

**For** $k = 1$**:** A family enters the "spread 1" bin via innovation. It leaves if it expands to spread 2.

$$\Delta m_{1}=m_{1}\left( t+1 \right)-m_{1}\left( t \right)=\mu-\left( 1-\mu\right)\frac{1\cdot m_{1}\left( t \right)}{t}.\quad$$

Eq S2

**2.2 Asymptotic Solution**

We seek an asymptotic scaling solution where the fraction of cluster with spread $k$ remains constant over time, i.e., $m_{k}\left( t \right)\approx M_{k}\cdot t$.

Specifically, $M_{k}$ denotes the **steady-state probability density** (or relative frequency) of BGC clusters occupying exactly $k$ clades. In this framework, while the absolute number of clusters $m_{k}\left( t \right)$ grows as the system expands over time $t$, their **relative proportion** $M_{k}$ remains invariant, capturing the time-independent structural signature of the evolutionary process.

From Eq S2:

$$M_{1}=\mu-\left( 1-\mu\right)M_{1}\Rightarrow M_{1}\left( 1+1-\mu\right)=\mu\Rightarrow M_{1}=\frac{\mu}{2-\mu}.$$

Eq S3

From Eq S1:

$$M_{k}=\left( 1-\mu\right)\left( k-1 \right)M_{k-1}-\left( 1-\mu\right)kM_{k},$$

$$M_{k}\left[ 1+\left( 1-\mu\right)k \right]=\left( 1-\mu\right)\left( k-1 \right)M_{k-1},$$

$$\frac{M_{k}}{M_{k-1}}=\frac{\left( 1-\mu\right)\left( k-1 \right)}{1+\left( 1-\mu\right)k}=\frac{k-1}{k+\frac{1}{1-\mu}}.$$

Eq S4

**2.3 Power-Law Asymptotics**

For large $k$, the ratio $\frac{M_{k}}{M_{k-1}}$ can be approximated using the property of the Beta function or the Gamma function:

$$M_{k}=M_{1}\frac{\Gamma\left( k \right) \Gamma!\left( 1+\frac{1}{1-\mu} \right)}{\Gamma!\left( k+1+\frac{1}{1-\mu} \right)}.$$

Eq S5

Applying the Stirling approximation $\frac{\Gamma\left( k \right)}{\Gamma\left( k+a \right)}\sim k^{-a}$, the distribution $M_{k}$ follows a power-law tail:

$$M_{k}\propto k^{-\alpha}\quad\text{where}\quad\alpha=1+\frac{1}{1-\mu}.$$

Eq S6

$\alpha$ is the exponent of the Probability Density Function (PDF).

**3. Relationship between Zipf Exponent (**$\boldsymbol{\beta}$**) and Innovation (**$\boldsymbol{\mu}$**)**

In our analysis, we visualized the **Rank-Abundance distribution**, where the spread $S$ for a specific cluster is plotted against its descending rank $R$. According to Zipf's Law:

$$S\propto R^{-\beta}$$

Eq S7

In the asymptotic continuous approximation, the relationship between the Zipf exponent $\beta$ and the PDF exponent $\alpha$ is given by:

$$\beta=\frac{1}{\alpha-1}$$

Eq S8

Substituting our derived $\alpha=1+\frac{1}{1-\mu}$ into this relation:

$$\beta=\frac{1}{\left( 1+\frac{1}{1-\mu} \right)-1}=\frac{1}{\frac{1}{1-\mu}}=1-\mu$$

Eq S9

**Conclusion**

For NRP-siderophores, the observed $\beta= 0.73$ yields an innovation rate of $\mu\approx0.27$. As taxonomic scale increases, the decreasing $\beta$ likely reflects an increasing effective innovation rate, as high-level clade barriers suppress the expansion efficiency of existing clusters.

### Supplementary Figures

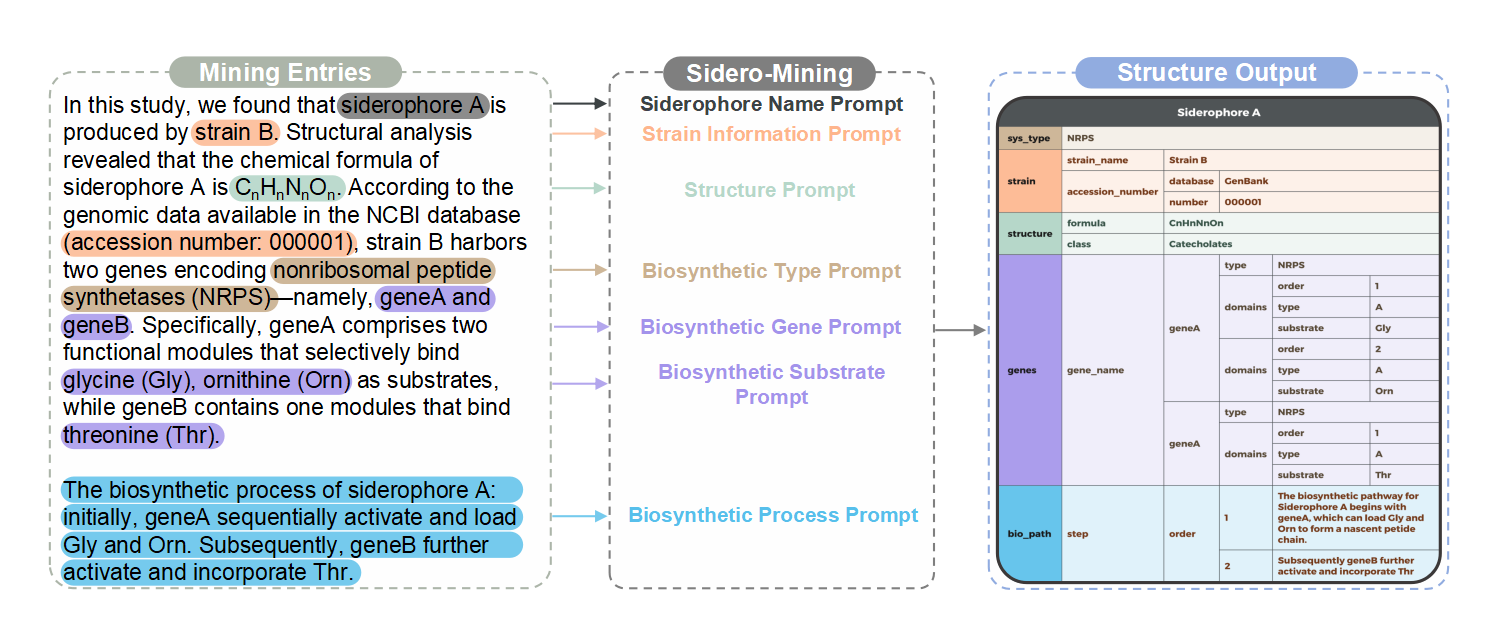

**Figure S1. Sidero-Mining Framework for Extraction and Structured Organization of Siderophore Information from Scientific Literature**

The figure illustrates the Sidero-Mining workflow for extracting siderophore information from research papers. Left panel ("Mining Entries") shows a sample text entry containing information about a hypothetical “siderophore A”’s production, chemical formula, and

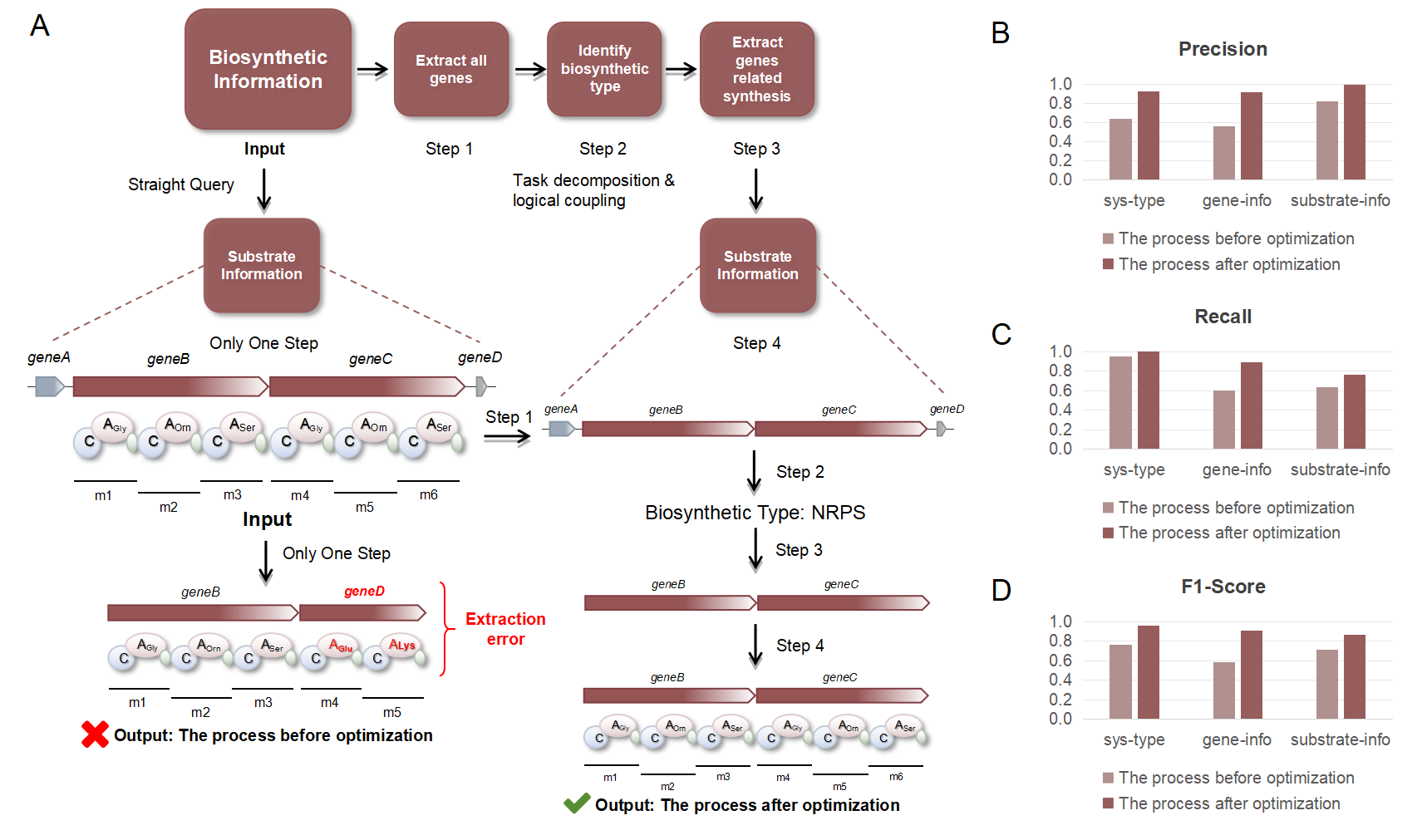

**Figure S2. Improving NRPS Substrate Extraction Through Task Decomposition in Sidero-Mining**

(A) Schematic comparison of traditional versus optimized substrate extraction workflows. The optimized process breaks down the complex task into four logical steps: extracting all siderophore-related genes (Step 1), determining synthesis type based on gene function (Step 2), identifying genes specifically related to synthesis (Step 3), and extracting substrate specificity information (Step 4). The diagram contrasts this with the traditional single-step approach, which often resulted in incomplete or inaccurate extraction.

(B-D) Performance comparison before and after workflow optimization across three key metrics: (B) Precision, (C) Recall, and (D) F1-Score. Results are shown for three extraction tasks: synthesis type identification (sys-type), gene information extraction (gene-info), and substrate information extraction (substrate-info). The optimized task decomposition approach shows substantial improvement across all metrics.

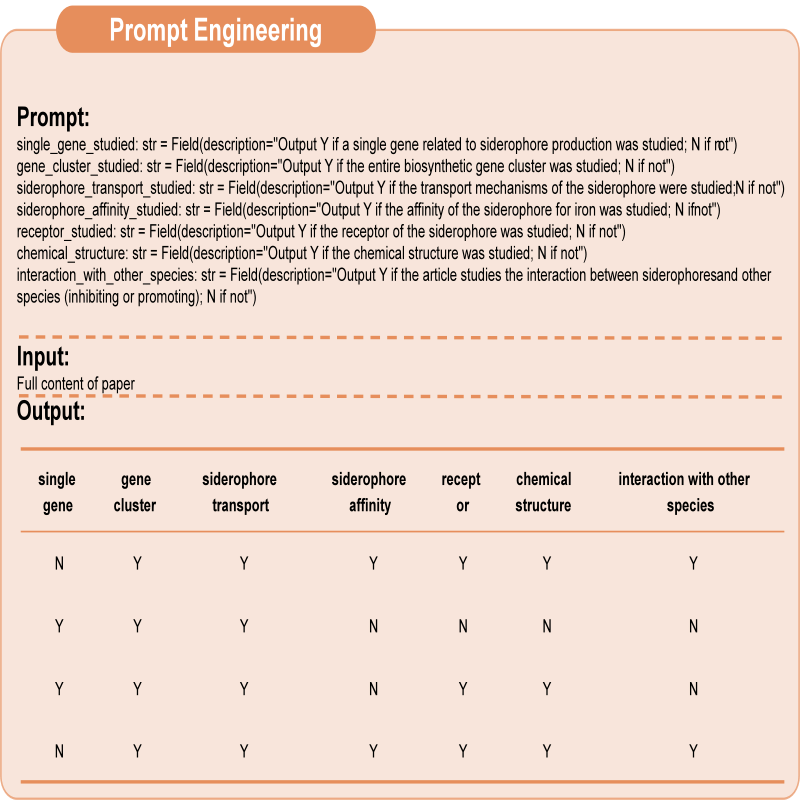

**Figure S3.** A schematic diagram of article screening. The full text of each article is input into GPT-4o-mini (gpt-4o-mini-2024-07-18), and seven preset prompts are applied—each targeting single-gene research, genomic research, siderophore transport, iron affinity determination, siderophore receptors, chemical structure research, and microbial interactions—to evaluate the article, with the model required to answer using “Y” or “N”.

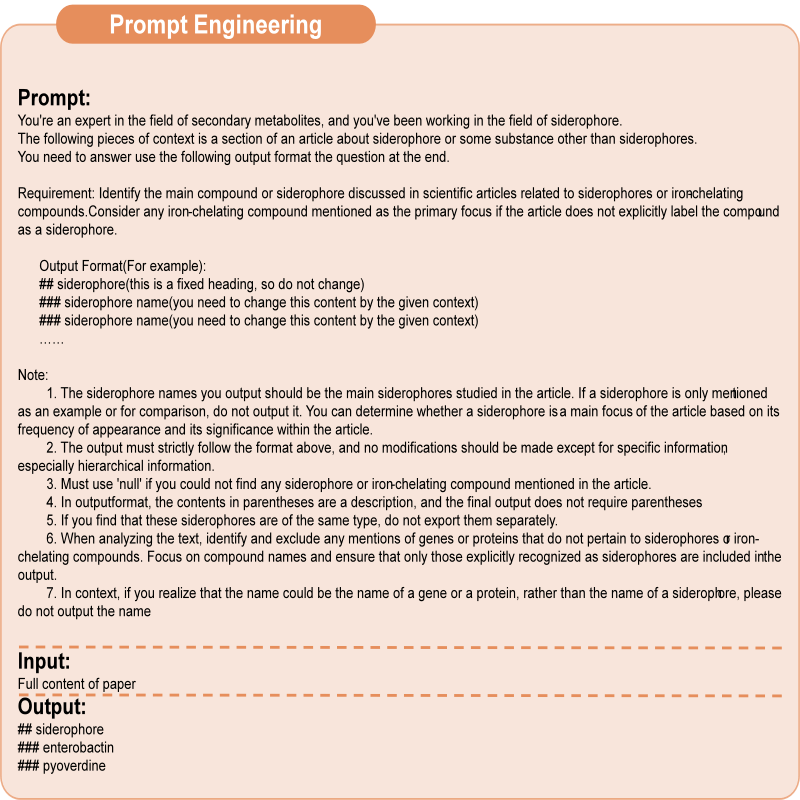

**Figure S4.** A schematic diagram for extracting siderophore names. After inputting the full text of the article and providing the model with the extraction requirements for siderophore names along with an output format template, the model returns the siderophore names as specified.

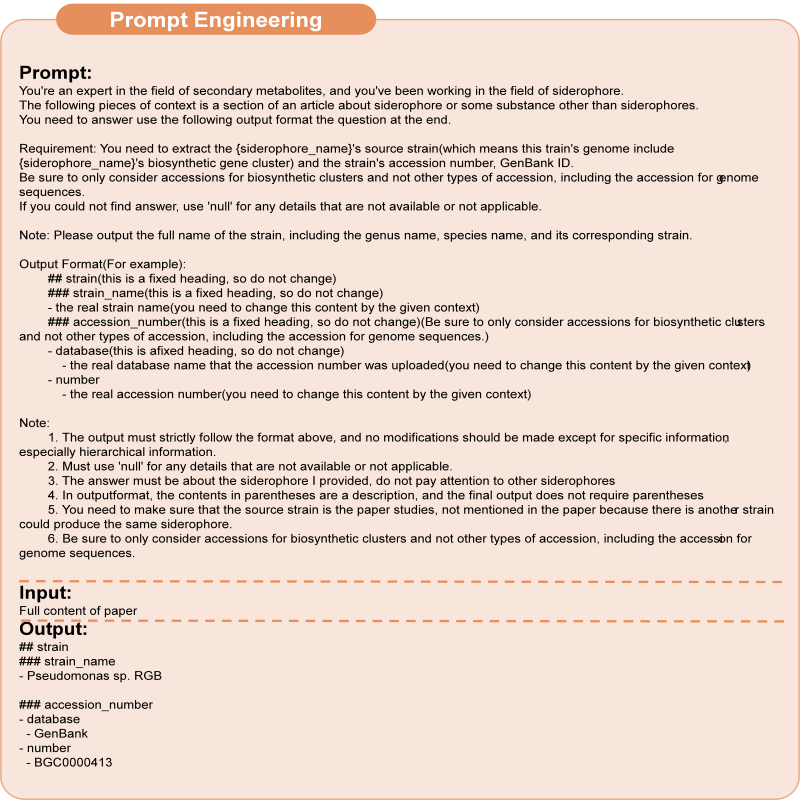

**Figure S5.** A schematic diagram for extracting siderophore-producing strain information. After inputting the full text of the article and providing the model with the extraction requirements and output format template, the model returns the relevant information for the target siderophore-producing strains as specified.

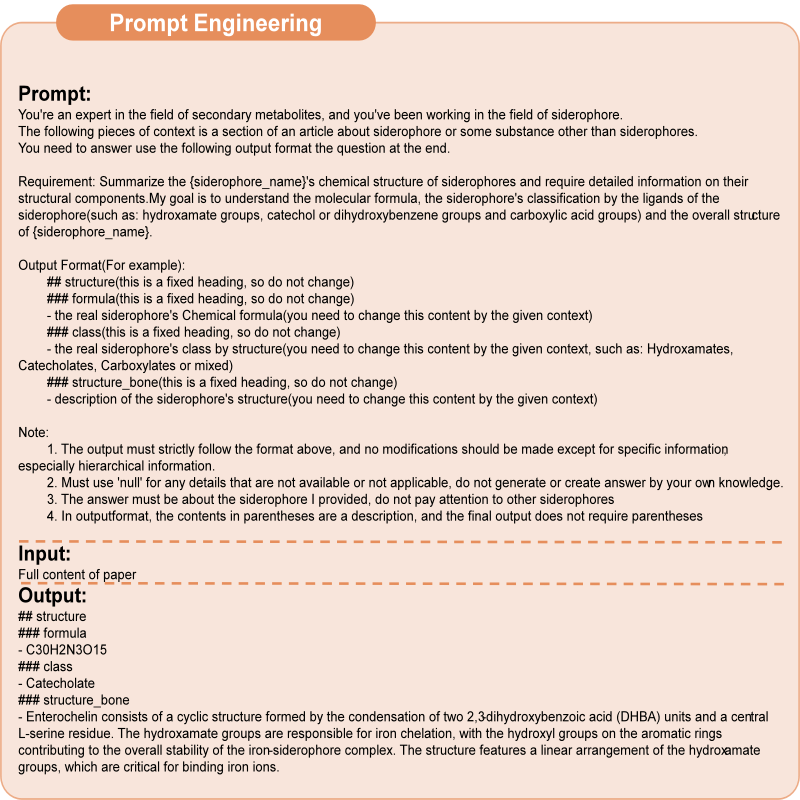

**Figure S6.** A schematic diagram for extracting siderophore chemical structure information. After inputting the full text of the article and providing the model with the extraction requirements and output format template, the model returns the relevant information for the target siderophore's chemical structure as specified.

Note: The article text must include the chemical formula of the siderophore for the model to extract the corresponding information.

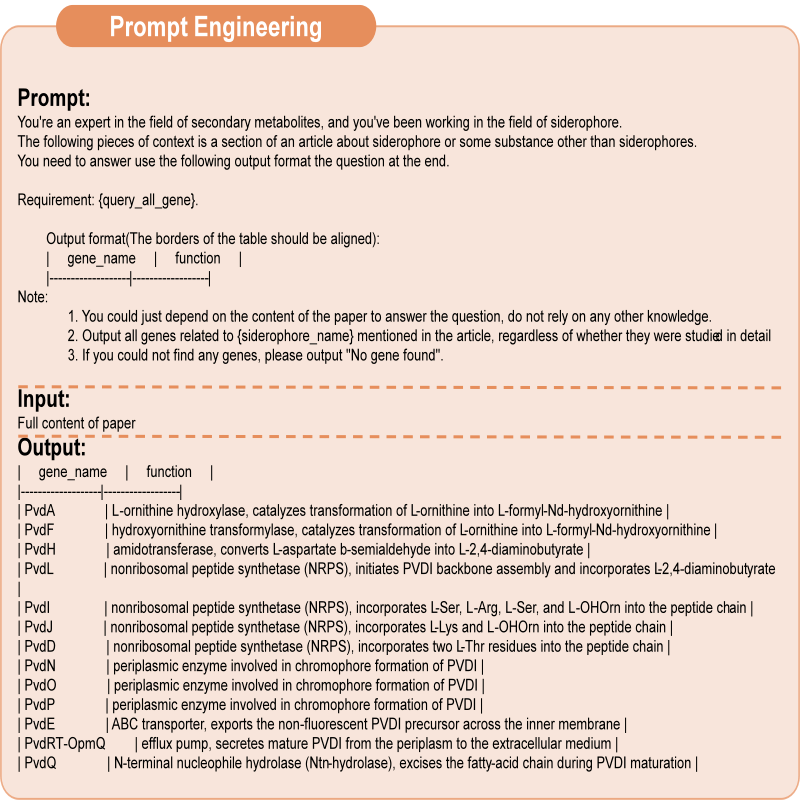
**Figure S7.** A schematic diagram for extracting all genes related to the target siderophore. After inputting the full text of the article and providing the model with the extraction requirements and output format template for all siderophore-related genes, the model returns the relevant gene information as specified.

Note: This part is an internal task within the Sidero-Mining workflow and is not part of the final answer. Therefore, it is not output in the standard Markdown format; instead, a text-based table using '|' and '-' is generated to serve as input for the next task.

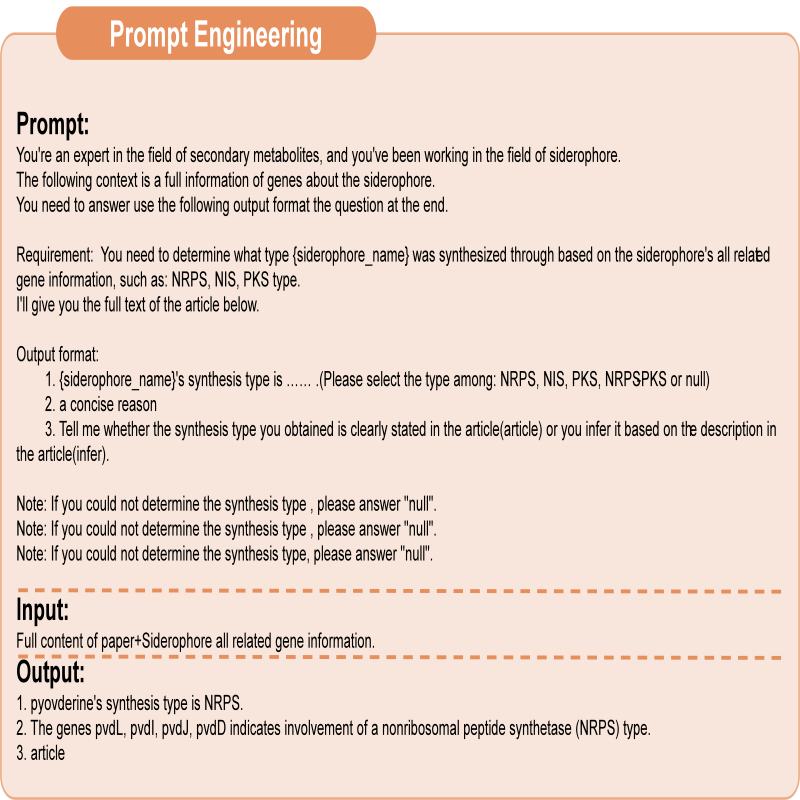

**Figure S8.** A schematic diagram for extracting the target siderophore synthesis type. By inputting the output from the previous step (Figure S5, which contains all genes related to the target siderophore) along with the full text of the article and providing the model with the extraction requirements for the siderophore synthesis type, the model returns the synthesis type information as specified. This includes the synthesis type, the rationale, and whether the answer was directly extracted from the article or inferred from its content.

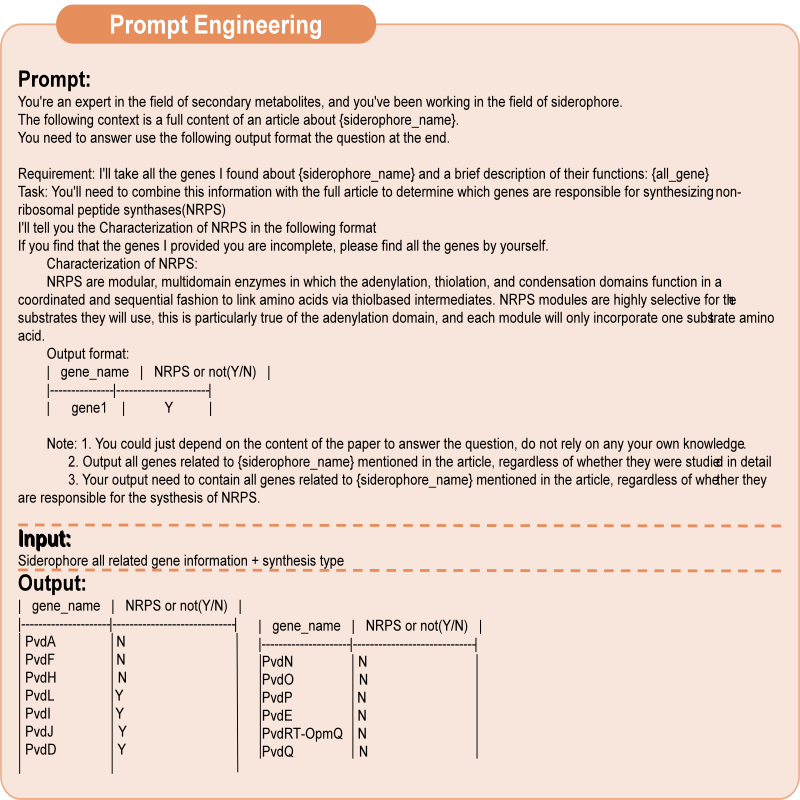

**Figure S9.** A schematic diagram for extracting the target siderophore synthesis genes. By inputting the complete set of genes related to the target siderophore (Figure S5), the siderophore synthesis type (Figure S6), and the task requirements into the model, the model returns the determination of all the target siderophore genes as specified.

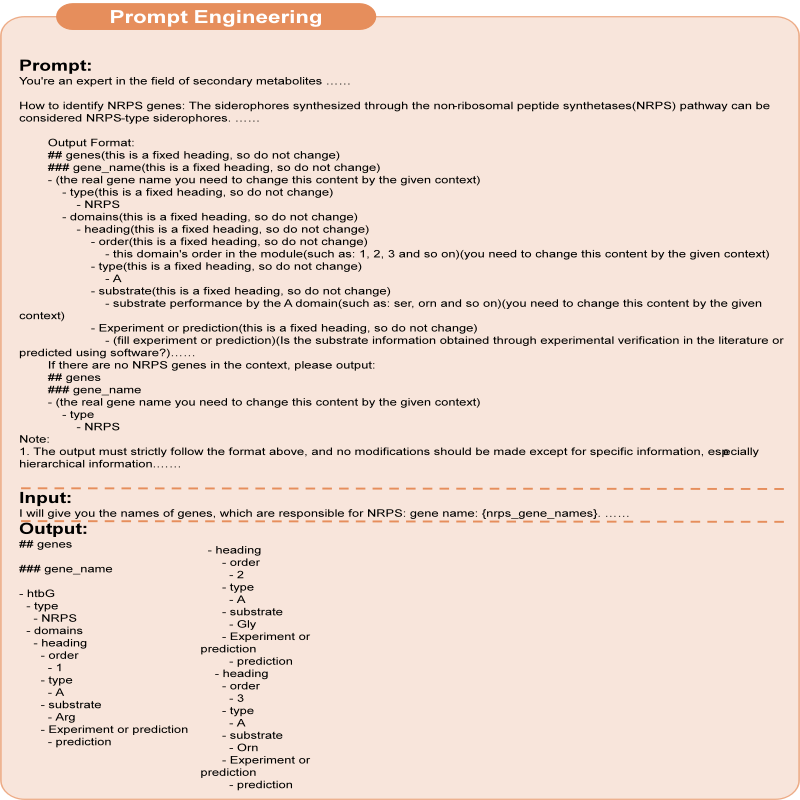

**Figure S10.** A schematic diagram for extracting substrate information for an NRPS siderophore. By inputting the siderophore synthesis genes (Figure S7) and the full text of the article, along with providing the extraction requirements and output format template for substrate information, the model returns the relevant substrate details for the target siderophore synthesis as specified.

Note: To ensure clear substrate information, the hierarchical structure of the output must be strictly defined.

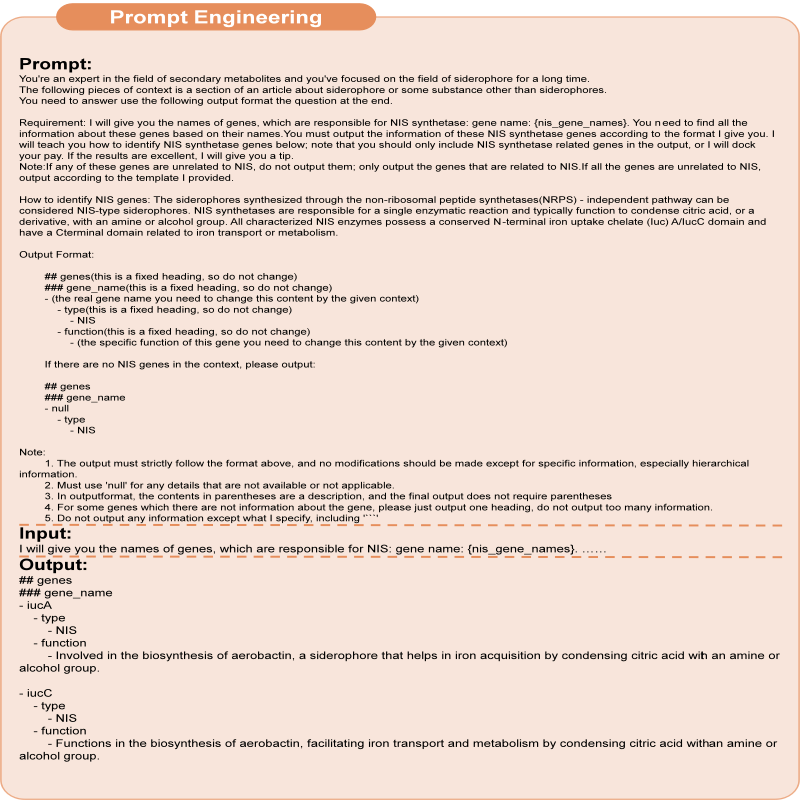

**Figure S11.** A schematic diagram for extracting synthesis information for an NIS siderophore. By inputting the siderophore synthesis genes (Figure S7) and the full text of the article, along with providing the extraction requirements and output format template for synthesis information, the model returns the relevant details for the target siderophore synthesis as specified.

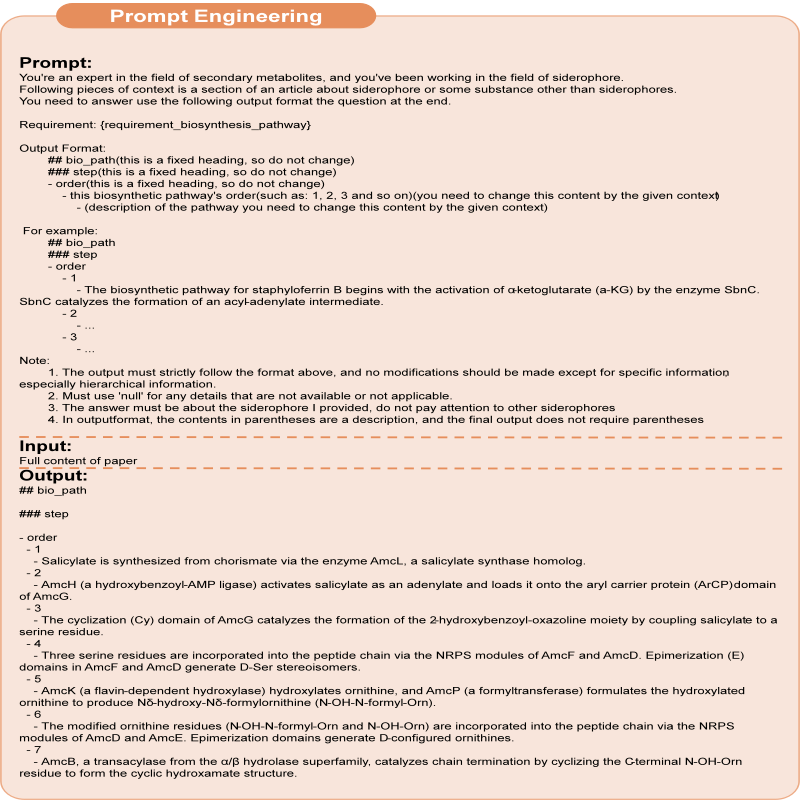

**Figure S12.** A schematic diagram for extracting the siderophore synthesis process. By inputting the full text of the article and providing the extraction requirements along with the output format template for the synthesis process, the model returns the relevant details of the target siderophore synthesis process as specified.

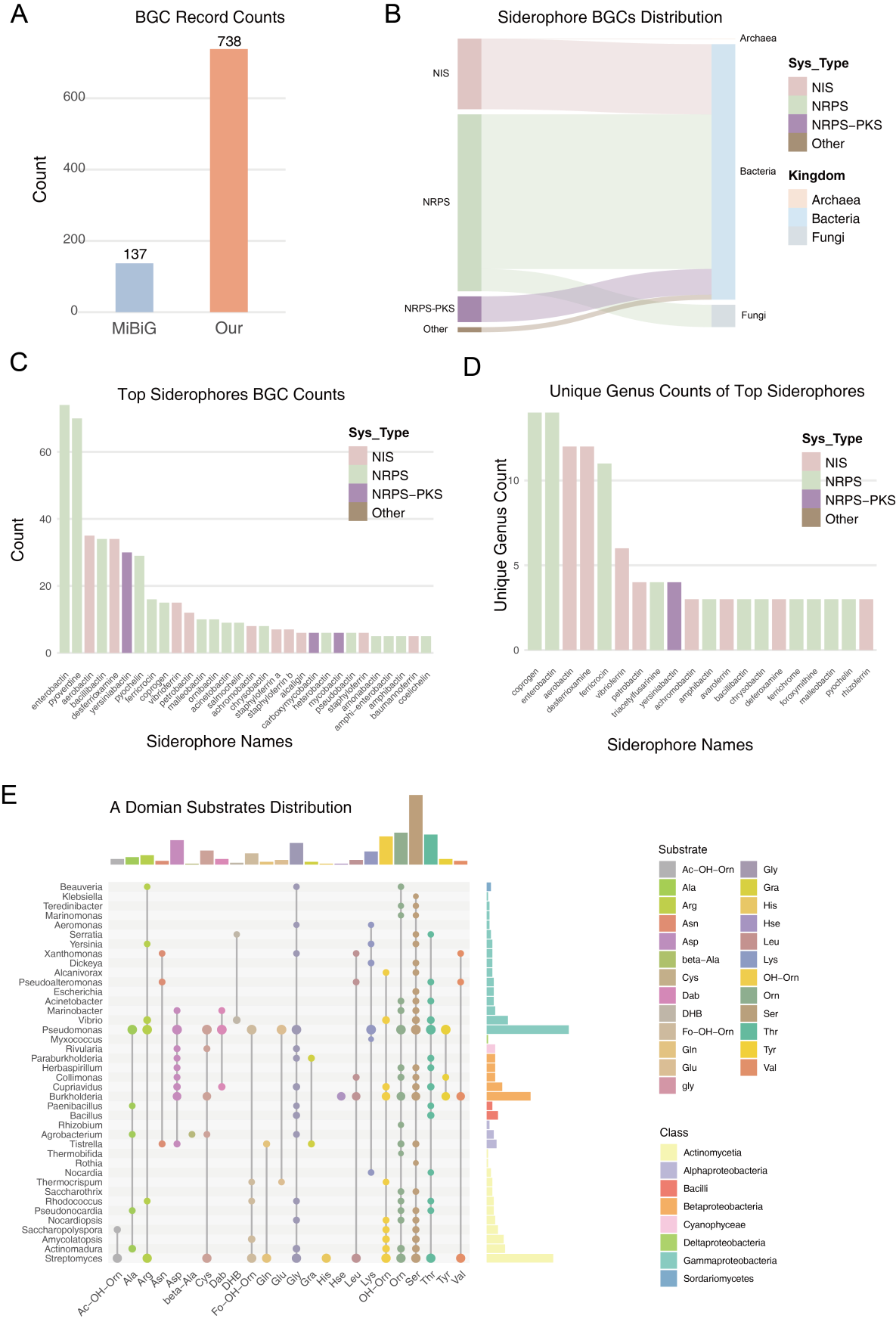

**Figure S13. A domain collection**

(E) The distribution patterns of A domain substrates and their respective source species. Substrates are differentiated by distinct colors, while species from various genera are color-coded according to their taxonomic class.

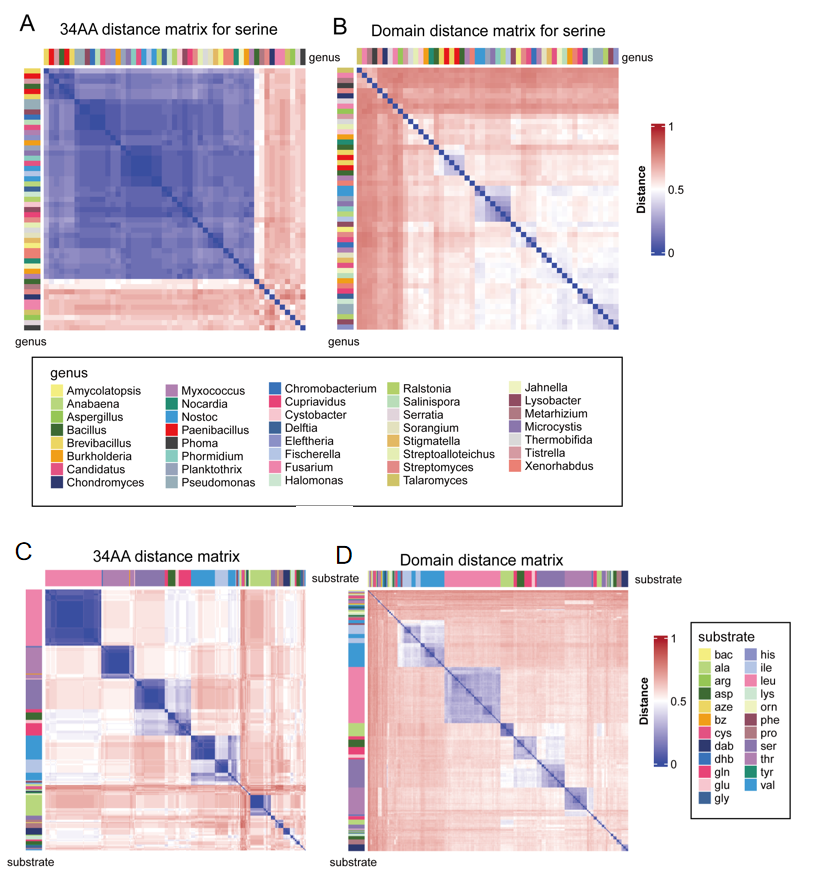

**Figure S14 Comparison of Substrate Differentiation Ability Between 34AA and Full-Length Domains**

(A-B) Distance matrices for serine-recognizing A domains based on 34AA and full-length sequences.

(C-D) Distance matrices for A domains of different substrates in Pseudomonas based on full-length and 34AA features.

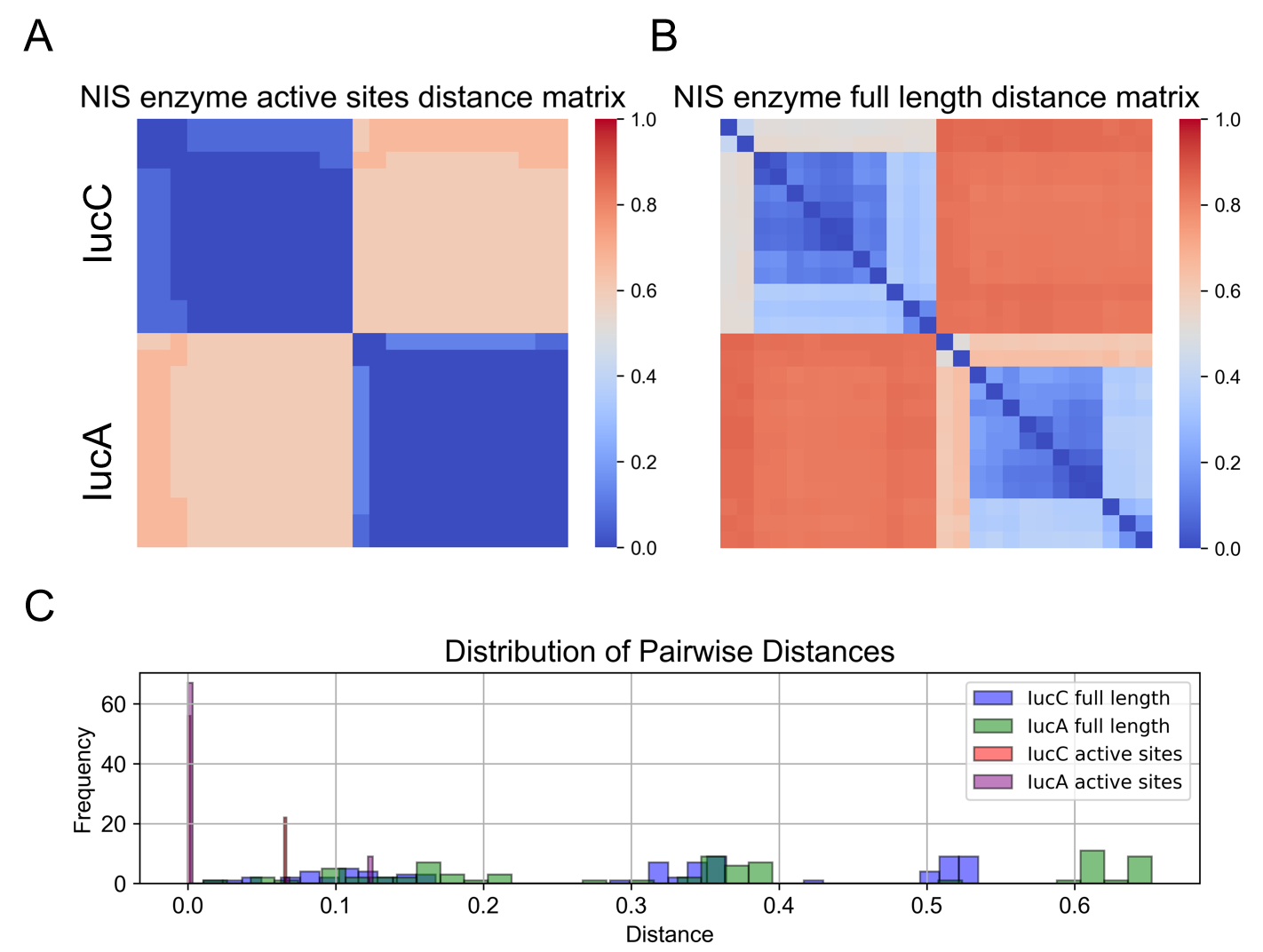

**Figure S15 Comparison of NIS Synthases in Aerobactin Across Genera: Active Site Features vs. Full-Length**

(A) Distance matrix for IucA and IucC based on active site features.

(B) Distance matrix for IucA and IucC based on full-length sequences.

(C) Pairwise distance distribution of the two synthases under both features.

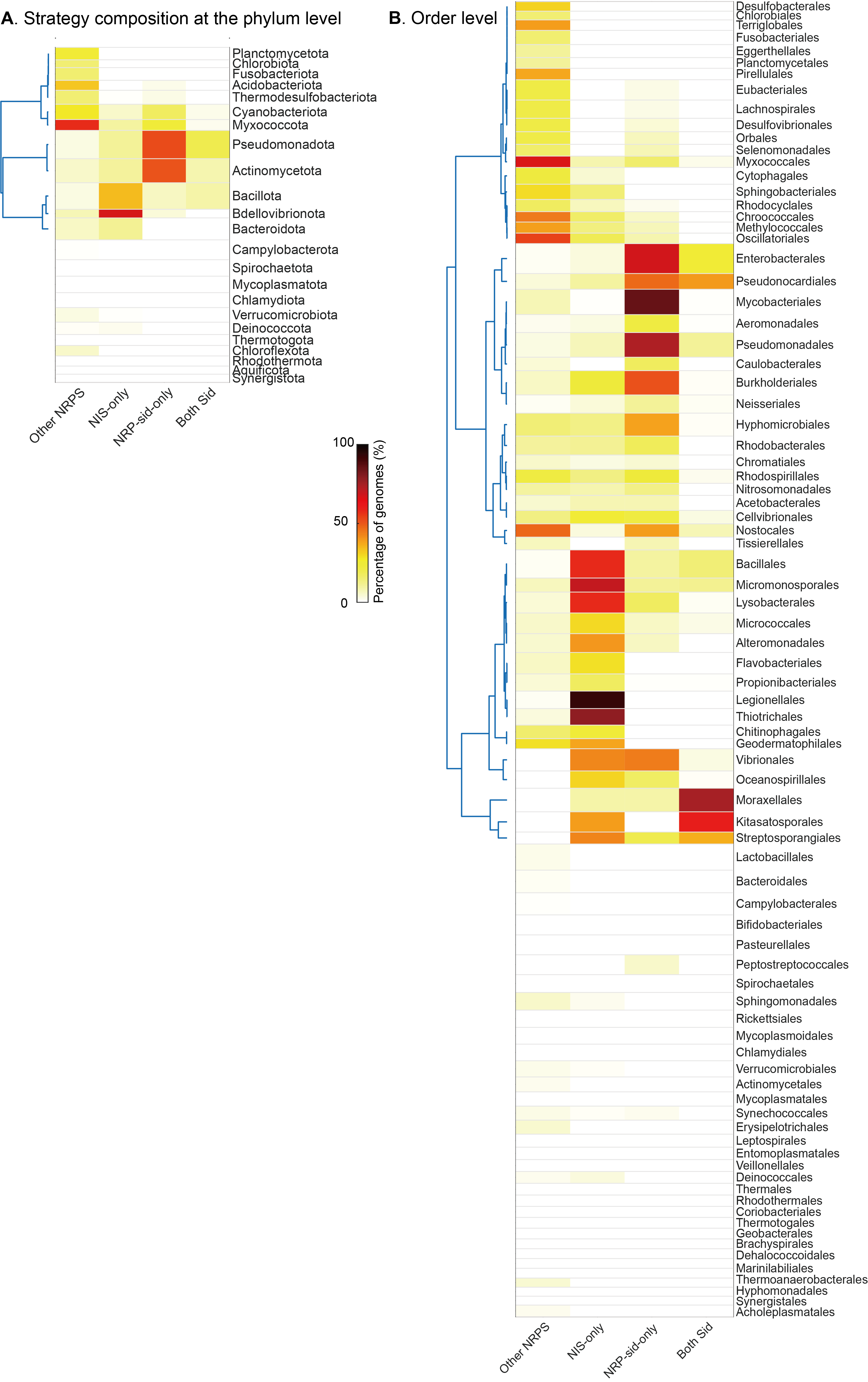

**Figure S16. Distribution of siderophore biosynthetic strategies across different clades.** The heatmap quantifies the percentage of genomes within each bacterial clades at the phylum level (A) or the order level (B), that adopts one of four mutually exclusive pathways: *Other NRPS* (possessing NRPS machinery but lacking dedicated siderophore BGCs); *NI-Sid only*; *NRP-Sid only*; and *Both Sid* (co-occurrence of both systems). Bacterial clades are vertically ordered by hierarchical clustering based on strategy profile similarity.

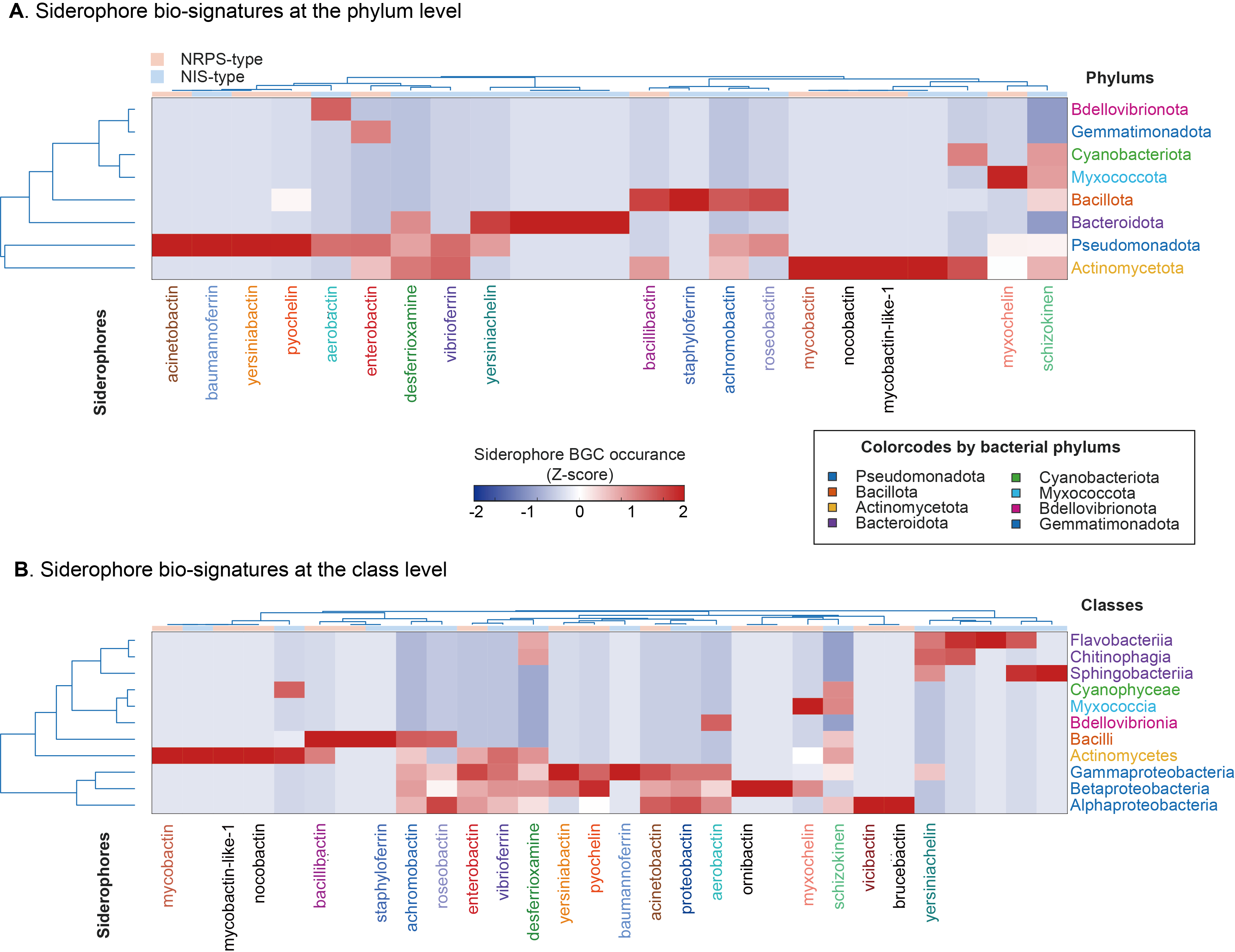

**Figure S17. Biosignature landscape of siderophore potential across bacterial clades.** A dual-hierarchical clustering heatmap illustrating the occurrence of distinct siderophore BGCs across varied orders. Color intensity denotes the Z-score normalized occurrence of each BGC (red: high; dark blue: absent/low). Rows represent bacterial clades at the phylum level (A) and the class level (B) , with labels color-coded by their parent class. Columns represent individual siderophore BGCs; labels for focal BGCs are colored consistently with plot B, non-focal BGCs with annotations are colored black, while unannotated BGCs are omitted for visual clarity. The top annotation band delineates the overarching biosynthetic logic: NRPS-type (peach) versus NIS-type (light blue). Clustering was performed using Ward’s linkage based on the Euclidean distance of prevalence profiles.

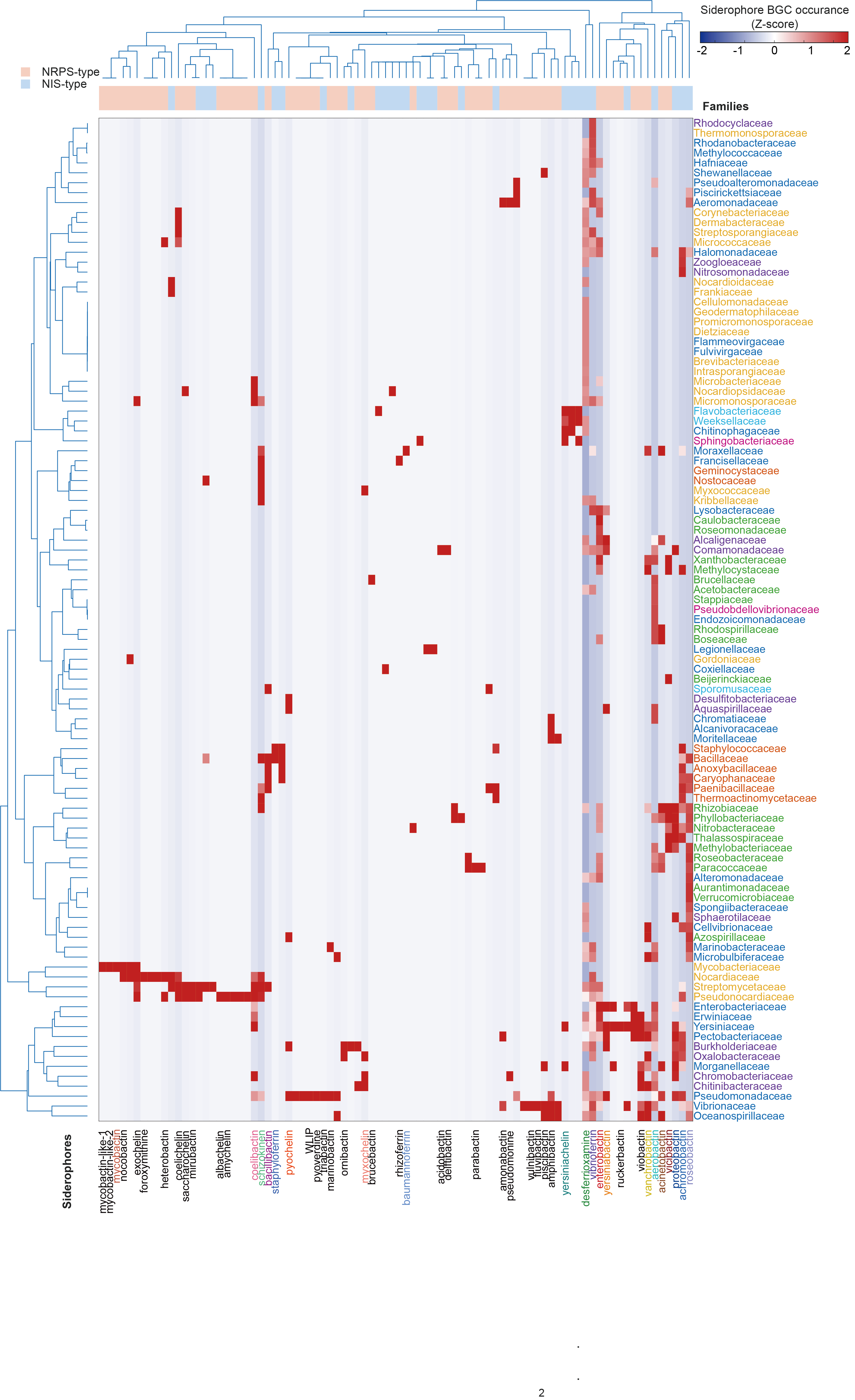

**Figure S18. Biosignature landscape of siderophore potential across bacterial families.** Same as Figure S_B, but the taxonomic clades are on the family level.

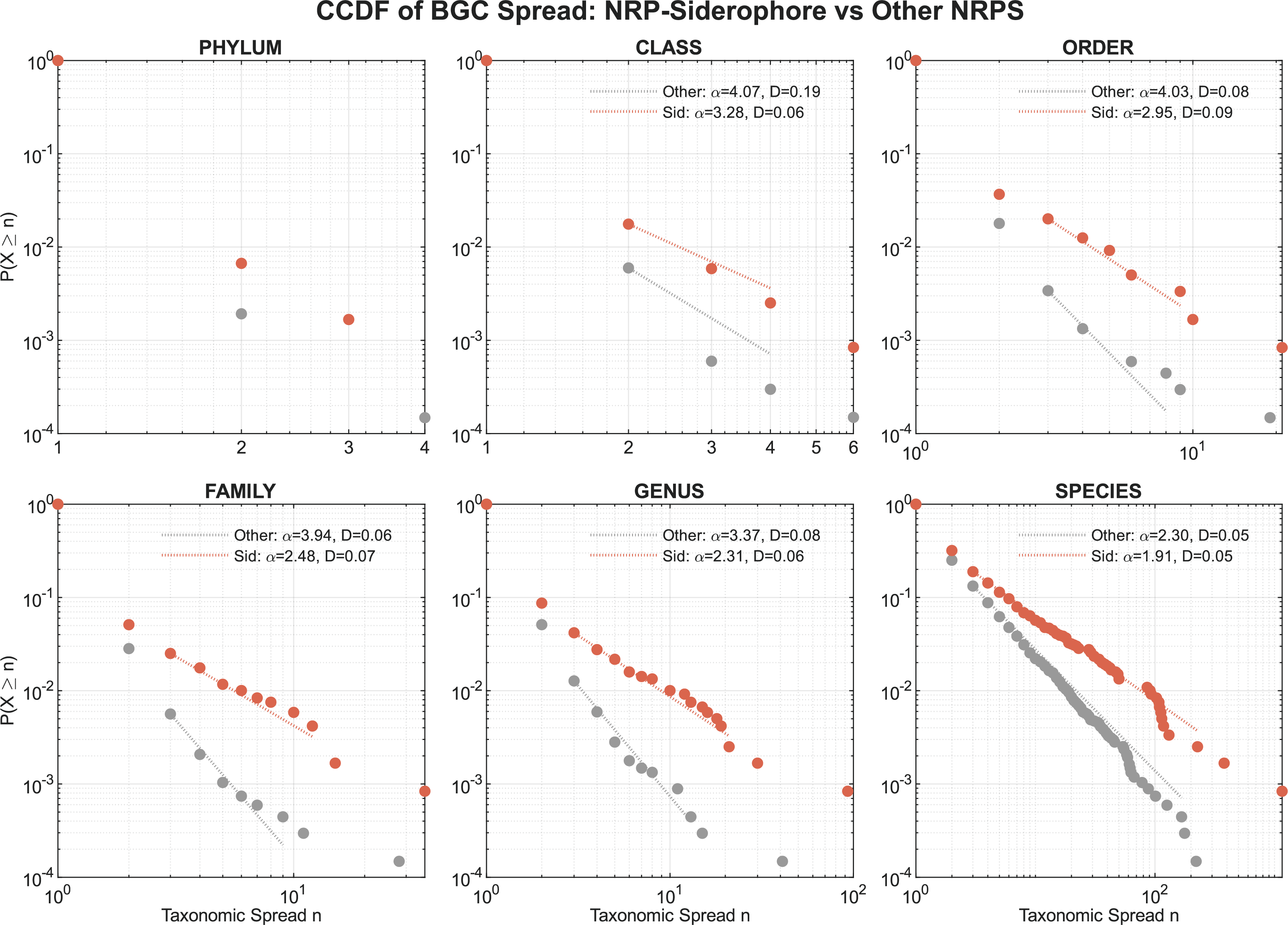

**Figure S19. Complementary Cumulative Distribution Function (CCDF) of BGC taxonomic spread**. Each panel compares the spread distribution of siderophore (termed "Sid“, salmon color ) versus non-siderophore NRPS (termed ”Other“, grey color) across six taxonomic levels. Circular markers represent empirical data, while dashed lines show the Maximum Likelihood Estimation (MLE) power-law fits. The calculated power-law scaling parameter ($\alpha$) and Kolmogorov-Smirnov (KS) statistic (D) are provided in the legends. The axes are shown in log-log scale, representing the probability$P\left( x\geq n \right)$ that a BGC is detected in at least $n$ distinct taxa at the corresponding level.

### Supplementary Tables

Table S1 Final Performance of Sidero-Mining on Downstream Tasks

| Mining Elements | Precision | Recall | F1-Score |
| --- | --- | --- | --- |
| siderophore-name | 0.9833 | 1.0000 | 0.9916 |
| species-name | 0.9833 | 1.0000 | 0.9916 |
| sys-type | 0.9259 | 1.0000 | 0.9615 |
| gene-info | 0.9189 | 0.8947 | 0.9067 |
| substrate-info | 1.0000 | 0.7667 | 0.8679 |

Table S2 NIS reference protein structure sites

| Enzyme | PDB id | Active sites |
| --- | --- | --- |
| DesD | 7tgm | 153, 156, 299, 300, 301, 303, 438, 442, 443, 479, 556, 558, 569, 570, 579 |
| IucA | 5jm8 | 147, 284, 285, 288, 29, 423, 425, 445, 447, 448, 450, 471, 479, 482, 483 |
| Sbnc | 7cbb | 147, 292, 297, 298, 422, 449, 468, 490, 552 |
| AsbB | 3to3 | 282, 308, 311, 313, 434, 459, 503, 567 |

Table S3 Phylogenetic decay of siderophore prevalence across taxonomic levels

| Taxonomic level | Encoding NRP-Siderophores (%) | Encoding NI-Siderophores (%) | Encoding either Siderophore (%) |
| --- | --- | --- | --- |
| Genome | 48.33% | 29.97% | 64.75% |
| Species | 24.76% | 23.10% | 39.30% |
| Genus | 16.18% | 17.70% | 26.43% |
| Family | 23.82% | 23.97% | 33.28% |
| Order | 25.10% | 25.49% | 34.51% |
| Class | 16.98% | 20.75% | 27.36% |
| Phylum | 20.41% | 20.41% | 26.53% |
